# Temporal changes in the gut microbiota in farmed Atlantic cod (*Gadus morhua*) outweigh the response to diet supplementation with macroalgae

**DOI:** 10.1101/2020.08.10.222604

**Authors:** C. Keating, M. Bolton-Warberg, J. Hinchcliffe, R. Davies, S. Whelan, A.H.L. Wan, R. D. Fitzgerald, S. J. Davies, U. Z. Ijaz, C. J. Smith

**Author notes:** These authors contributed equally (C. Keating, M. Bolton-Warberg). **Correspondence:** Joint corresponding authors.

## Abstract

**Background:** Aquaculture successfully meets global food demands for many fish species. However, aquaculture production of Atlantic cod (*Gadus morhua*) is modest in comparison to market demand. For cod farming to be a viable economic venture specific challenges on how to increase growth, health and farming productivity need to be addressed. Feed ingredients play a key role here. Macroalgae (seaweeds) have been suggested as a functional feed supplement with both health and economic benefits for terrestrial farmed animals and fish. The impact of such dietary supplements to cod gut integrity and microbiota, which contribute to overall fish robustness is unknown. The objective of this study was to supplement the diet of juvenile Atlantic cod with macroalgae and determine the impacts on fish condition and growth, gut morphology and hindgut microbiota composition (16S rRNA amplicon sequencing). Fish were fed one of three diets: control (no macroalgal inclusion), 10% inclusion of either egg wrack (*Ascophyllum nodosum*) or sea lettuce (*Ulva rigida*) macroalgae in a 12-week trial.

**Results:** The results demonstrated there was no significant difference in fish condition, gut morphology or hindgut microbiota between the *U. rigida* supplemented fish group and the control group at any time-point. This contrasts with the *A. nodosum* treatment. Fish within this group were further categorised as either ‘Normal’ or ‘Lower Growth’. ‘Lower Growth’ individuals found the diet unpalatable resulting in reduced weight and condition factor combined with an altered gut morphology and microbiome relative to the other treatments. Excluding this group, our results show that the hindgut microbiota was largely driven by temporal pressures with the microbial communities becoming more similar over time irrespective of dietary treatment. The core microbiome at the final time-point consisted of the orders *Vibrionales* (*Vibrio* and *Photobacterium*), *Bacteroidales* (*Bacteroidetes* and *Macellibacteroides*) and *Clostridiales* (*Lachnoclostridium*).

**Conclusions:** Our study indicates that *U. rigida* macroalgae can be supplemented at 10% inclusion levels in the diet of juvenile farmed Atlantic cod without any impact on fish condition or hindgut microbial community structure. We also conclude that 10% dietary inclusion of *A. nodosum* is not a suitable feed supplement in a farmed cod diet.

## Background

Sustainable food production is gaining increased attention from consumers, farmers, and policymakers. This is spurred on by concerns on the impact of climate change, consumer awareness, and the growing global food demand [1]–[3]. Currently, 151 million tonnes of fish are produced annually for human consumption [4]. Wild fish stocks cannot sustain further increases in demand. Several wild fish populations have collapsed or are in danger of critical decline from over-fishing and habitat damage. A significant depletion in 4 European cod stocks (Kattegat, North Sea, West of Scotland and Irish Sea) by the early 2000s led to wild stock enhancement initiatives, such as the European Union’s Cod Recovery Plan (EC No 1342/2008; [5], and ultimately included enforced catch quotas. However, despite their initial recovery, The International Council for the Exploration of the Sea (ICES) have advised that the North Sea, Celtic Sea, Irish Sea and West of Scotland cod stocks are in decline once more [6]–[9]. The cause of this decline is unclear, with climate change, destruction of nearshore cod nurseries by trawling, grey seal predation and fisheries management being implicated [10]–[12]. The precarious condition of these wild cod stocks means this is simply not a sustainable option to meet consumer demand for this fish.

Commercial aquaculture has successfully met the market demand for a range of fish species, including Atlantic salmon (*Salmo salar*), rainbow trout (*Oncorhynchus mykiss*), gilt-head seabream (*Sparus aurata*), and seabass (*Dicentrarchus labrax*), thereby alleviating demand on wild resources [4]. Despite the popularity of cod as a food fish, aquaculture production of farmed cod is a business that has waxed and waned since the 1970’s. Currently, only 2.5% of the total Atlantic cod produced for food is through aquaculture [3]–[4]. The economic viability of a commercial cod farming operation presents a number of challenges including a consistent high-quality broodstock, reducing high mortality, disease susceptibility, market pricing and high costs associated with aquafeeds [13]–[15]. Developing novel feed ingredients and alternatives to marine by-products in formulated diets not only could improve the economic viability of commercial cod farming but could also have implications in the sustainability status of cod farming and consumer perception of farmed cod.

One potential solution is to supplement the diet of farmed cod with high-quality functional ingredients that provide health benefits, nutrition and are environmentally sustainable and low cost. The use of macroalgae (also known as seaweeds) as a potential feed supplement could further this sustainability agenda. The biomass can often be grown or sustainably harvested at quantities that would meet commercial aquafeed demand. Dietary supplementation of macroalgae has been trialled in many farmed fish species including Atlantic salmon (*S. salar*, [16], rainbow trout (*O. mykiss*, [17], European seabass (*D. labrax*, [18], carp (*C. carpio*, [19]), and tilapia (*O. niloticus*, [20]. Some studies have advocated that macroalgae could partially replace protein constituents in formulated diets [21], [22]. However, the lower algal protein content (up to 27%) in comparison to highly proteinaceous fishmeal (e.g. LT94 fish meal, ~ 67 %) and plant by-products (e.g. soy protein concentrates, ~76%) typically used in formulated diets, indicate it would be better to exploit algal meal as a functional feed supplement instead [23]. As a functional feed supplement macroalgae can be used to supply trace metals, carotenoids, long-chain omega-3 polyunsaturated fatty acids, and vitamins that potentially benefit the growing farmed fish [24]. In general, the inclusion of macroalgae into formulated fish diets without causing decreased growth performance is approximately 10% [23]. However, the dietary macroalgae inclusion level that fish can tolerate is dependent on the algal species and fish species, i.e. carnivorous versus omnivorous fish, brown versus green macroalgae or indeed morphological differences in the fish gastrointestinal tract [25].

There is a wealth of evidence outside of fish research showing that the gut microbiota is intrinsically linked to digestive function, metabolism, immune support and general health as shown in dogs [26] and rats [27] for example. For fish gut microbiota these links are less clearly defined, though this is the focus of intensive research efforts in Atlantic salmon (*S. salar*) [28], [29]. We know that the digestibility and metabolism of a substrate is impacted by the gut microbiome [28], [30]. Fish gut microbiota produce exogenous digestive enzymes (e.g. lipases, amylases and proteases), which can significantly influence nutrient digestibility and uptake and renewal of the gut epithelial layer [30]. Research has also indicated that dietary amendment can cause changes in the gut microbiome which may or may not be associated with physiological changes to the fish [31]–[33]. Algae can possess unique taxon specific polysaccharides known as phycolloids (e.g. alginic acid and fucoidan), which have been shown using in vitro studies to exhibit antiviral, immunostimulatory, anticoagulant, and antioxidant bioactivity [34], [35]. In addition, complex components of dietary macroalgae such as cellulose, hemicellulose, and lignin, could alter the gut microbiome in the fish by providing a substrate source to hydrolytic bacteria [36]. It, therefore, follows that any manipulation of farmed cod diets with macroalgae could also impact the gut microbiome, which in turn may have yet to be understood implications for fish welfare and growth. Indeed, Gupta et al. [37] observed with prebiotic supplements containing alginate (extracted from *Laminaria* sp. of brown seaweed) corresponded to distinct microbiome changes in Atlantic salmon. Their study, however, did not comment on the associated health or growth impacts. Thus, a complementary approach combining fish growth, internal physiology, and the gut microbiome is more appropriate as microbiome changes alone are not indicative of condition or response to new feed supplements.

To further develop Atlantic cod aquaculture and to address the food sustainability agenda, this study aimed to evaluate if the diet of juvenile Atlantic cod could be supplemented with macroalgae and to determine the effects on fish growth, condition, gut morphology and gut microbiome structure. Two species of individual macroalgae dietary supplements were compared - egg wrack *Ascophyllum nodosum* (brown) and sea lettuce *Ulva rigida* (green), alongside a non-amended fishmeal diet. *A. nodosum* is a common algae species in northern temperate Atlantic waters and is harvested from the wild for commercial alginate (phycocolloid) production and farm animal feeds. *U. rigida* is ubiquitous and can be a nuisance species, proliferating in large quantities resulting in green tides. It is this significant algal biomass availability in the wild that would provide the upscale requirement in commercial aquafeed production [23]. The hypothesis was that Atlantic cod growth and condition would not be negatively impacted by macroalgae inclusion, but that the addition of the macroalgae would lead to changes in the gut microbiome compared to the control diet. To the author’s knowledge, this is the first study to examine the impact of dietary macroalgae supplementation on the gut integrity and hindgut microbiome in juvenile Atlantic cod.

## Methods

### Feed Formulation and Production

Macroalgae used in the feed trial were harvested from Muigh Inis, Co. Galway, Ireland (*Ascophyllum nodosum*) and Harbour View Bay, Co. Cork, Ireland (*Ulva rigida*, green tide, [38]). The biomass was washed with freshwater to remove debris and dried for 48 hrs at 40°C in a dehumidifying cabinet. The resulting dried biomass was milled using a hammer mill (Timatic, Spello, Italy) and sieved to a particle size of <0.8 mm. The proximate composition profile of the *U*. *rigida* biomass was 17.59 ± 0.09% protein, 0.48 ± 004% lipid, 22.53 ± 0.27% ash, and 10.08 ± MJ kg^-1^ energy. While, *A. nodosum* had 7.01 ± 0.10% protein, 0.94 ± 0.08% lipid, 24.83 ± 0.50% ash, and 11.77 ± 0.11 MJ kg^-1^ energy. The inclusion of macroalgae into the test diets was predominately at the expense of fish meal and potato starch. Diets were formulated to be nutritionally balanced and were iso-nitrogenous (50%), iso-lipidic (20%) and iso-energetic (20 MJ kg^-1^). Three diets (nominally CTRL, ULVA, ASCO, Table 1) were produced in-house through cold extrusion as described in Wan et al. [16]. Proximate composition of the finished diets was analysed to confirm diet quality [39]. Quantification of protein was carried out by Kjeldahl procedure (DT220 and Kjeltec 8200, Foss A/S, Hillerød, Denmark) using x6.25 conversion factor and lipid level was determined through Soxhlet extraction using petroleum ether (ST243, Foss A/S, Hillerød, Denmark). Ash content was measured through incineration of samples at 550 °C for 16 hrs. In addition, energy determination was carried out through bomb calorimetry (6200, Parr Instruments, Moline, Illinois, USA).

### Experimental Fish, Design and Fish Physical Condition

A total of 540 juvenile hatchery-reared Atlantic cod (*G. morhua*, the first-generation offspring of wild Celtic sea broodstock sourced in 2013) were employed in the feed trial. The stock fish of juvenile Atlantic cod (*Gadus morhua*) used were sampled prior to starting the experiment (Week 0). These fish had been fed a commercial fish pellet diet (Amber Neptun for marine fish, Skretting, Stavanger, Norway). Fish were hand-graded (123 ± 1 g) and randomly assigned to one of nine tanks (1200 L, *n*=60 per tank, 3 replicate tanks per diet). The research tanks were fed with flow-through filtered ambient seawater (13.0 ± 1.5°C) and were aerated to maintain a dissolved oxygen level of > 6 mg L^-1^. A photoperiod of 8:16 light:dark was employed throughout the experiment.

Fish were acclimated for one week during which all tanks were fed a control diet (Table 1; CTRL) before starting the experimental diets. Following this acclimation period, each tank was randomly assigned to one of three diets: Control (CTRL), 10% supplement of *Ulva rigida* (ULVA) and 10% supplement of *Ascophyllum nodosum* (ASCO). Fish were hand-fed three times daily (evenly spaced throughout the day). The feeding rate was ~1% of the total tank body weight and was adjusted every two weeks to account for mortality, temperature and growth using standard cod growth models [40], [41]. Briefly, as is standard practice, fish were starved for 24 hrs before sampling, for total body weight (g) and total length (cm), of all individuals per experimental tank to calculate tank population growth rate. Fish were sampled for tank population data once monthly from the start of the acclimation period (i.e. from Week 0). Any mortalities were removed daily and recorded.

**Table 1.** The experimental diet formulations and their proximate composition (%, dry weight).

| Diet formulation | Diet |  |  |
| --- | --- | --- | --- |
|  | CTRL | ULVA | ASCO |
| LT94 Fish meal<sup>a</sup> | 66.00 | 64.94 | 62.44 |
| Fish oil <sup>a</sup> | 12.74 | 12.75 | 12.90 |
| *Ascophyllum nodosum* | - | - | **10.00** |
| *Ulva rigida* | - | **10.00** | - |
| Wheat gluten<sup>b</sup> | 9.00 | 9.00 | 9.00 |
| Potato starch<sup>b</sup> | 10.10 | 1.18 | 2.53 |
| Vitamin & mineral premix<sup>c</sup> | 2.00 | 2.00 | 2.00 |
| Antioxidant<sup>d</sup> | 0.02 | 0.02 | 0.02 |
| **Proximate Composition\*** |  |  |  |
| Protein | 50.65 | 50.76 | 50.15 |
| Lipid | 20.10 | 20.79 | 20.82 |
| Ash | 15.53 | 18.05 | 17.58 |
| Energy; MJ kg<sup>-1</sup> | 20.12 | 20.12 | 19.96 |

\*n=3.<sup>a</sup> United fish industries Ltd, Grimsby, UK. <sup>b</sup> Gemcom, London, UK <sup>c</sup> Premier nutrition products Ltd., Rugley, UK. (Manufacturer's analysis: Ca 12.09%; ash 78.71 %; Na 8.86 %; vitamin A 1.00 µg kg<sup>-1</sup>; vitamin D<sub>3</sub> 0.10 %, vitamin E 7.00 g kg<sup>-1</sup>; Cu 250 mg kg<sup>-1</sup>; Mg 15.6 g kg<sup>-1</sup>; P 5.2 g kg<sup>-1</sup>) <sup>d</sup> Barox plus liquid, Kemin Europa N.V., Herentals, Belgium.

From Week 8 on, it was noted via visual distinction (Supplementary Information: Figure S1) that there were two subsets of fish within the *A. nodosum* supplemented diet: those that found the diet palatable and those that did not. This was confirmed by the presence of uneaten feed pellets in ASCO tanks following feeding events (not visible in tanks fed with CTRL or ULVA diets). Individuals that found the diet unpalatable had a markedly reduced size. These groups were referred to as ASCO_N (N – Normal) and ASCO_LG (LG – Lower Growth).

### Sample Collection

For both histology and microbiome analysis, sacrificial sampling was necessary. Fish were humanely euthanised using an overdose of tricaine methanesulphonate solution (MS222, Pharmaq, Overhalla, Norway), which was followed by the destruction of the brain to confirm death (EU Directive 2010/63/EU). The number of fish used for sacrificial sampling was kept to a minimum while remaining statistically sound. Separate individuals were necessary for histological and hindgut microbiome measurements as the former required a 24 hrs starvation period while the latter required gut content. For all fish sampled, the total weight (g) and total length (cm) of each fish were recorded.

*Note:* Extra samples were taken from the ASCO treatment during the Week 8 and 12 sampling periods, with individuals classified as ‘ASCO_N’ or ‘ASCO_LG’ based initially on a visual assessment (i.e. slimmer body shape) and confirmed by calculation of condition factor. For histology analysis, three fish were taken per treatment (CTRL, ULVA, ASCO_N and ASCO_LG) at Week 8 and Week 12. For microbiome analysis, eight fish were taken at Week 0, 28 fish at Week 8 (6 CTRL, 9 ULVA, 8 ASCO_N, 5 ASCO_LG) and 31 fish at Week 12 (8 CTRL, 8 ULVA, 8 ASCO_N, 7 ASCO_LG). Variability in microbiome sample numbers was due to some fish not having sufficient gut content for analysis, resulting in additional fish being sampled.

#### Gut Morphology Analysis

Tissue samples for light microscopy (LM) and scanning electron microscopy (SEM) were taken in Weeks 8 and 12. Whole intestines (from the stomach to the anus) were dissected from each fish and the gastrointestinal (GI) tract unfolded and removed. The GI tract was subsequently flushed with phosphate buffer saline (PBS, pH 7.3) to remove gut digesta and mucosal layer. The total length of the intestine was measured and to standardise sampling, the hindgut defined as 10-15% of total gut length. The hindgut was subsequently fixed in 10% neutral buffered marine formalin at room temperature for 24 hrs. Samples were subjected to serial dehydration steps in alcohol (100, 90, 70, 50, and 30%) before equilibration in xylene in a tissue processor. Samples were then embedded in paraffin wax according to standard histological procedures [42]. Transverse sections (6 µm) were sectioned and mounted on silane-covered (TESPA, 3-aminopropyl-triethoxysilane, Sigma-Aldrich, St. Louis, Missouri, USA) glass slides. Slides were stained with haematoxylin and eosin and mounted for light microscopy analysis. Slides were examined by light microscopy using an Olympus Vanox-T microscope and images were taken with a digital camera (Olympus camedia C-2020 Z).

For scanning electron microscopy analysis (SEM), hindgut samples were dipped in 1% 5-carboxymethyl-L-cysteine (Sigma-Aldrich, St. Louis, Missouri, USA) PBS solution for 30 seconds to remove surface mucus and digesta. Samples were fixed in 2.5 % glutaraldehyde (pH 7.2) for 24 hrs and were dehydrated gradually in serial dilutions of 30, 50, 70, 90 and 100 % ethanol for critical point drying (K850, Emitech Ltd, Ashford, Kent, UK). The resulting samples were mounted on stubs with the use of agar silver paint and gold sputter coated (K550, Emitech Ltd, Ashford, Kent, UK). Images of the microvilli density were taken using a low vacuum scanning electron microscope (5600, Jeol, Tokyo, Japan).

#### Gut Microbiome Analysis

##### Collection of Hindgut Content

The undersides of the fish were swabbed with ethanol before dissection and removal of the gut. The length of the digestive tract was recorded (cm). The hindgut was defined as 10-15% of the total gut length and dissected. The gut contents of the hindgut were collected, flash frozen and stored at −80°C for subsequent molecular analysis.

##### Sample DNA Extraction and 16S rRNA Amplicon Sequencing

DNA extractions were carried out as described using a modified Griffiths et al. (2000) procedure [43], [44]. Briefly, 250 μL TE buffer was added to the contents of the hindgut (digesta) while still frozen, mixed, and added to Lysing Matrix E tubes (MP Biomedical, Illkirch-Graffenstaden, France) containing 500 μL cetyl trimethylammonium bromide (CTAB) buffer and 250 μL of phenol-chloroform-isoamyl alcohol (25:24:1; pH 8). The mixture containing 0.25 g of digesta content was lysed by bead beating for 10 min at 3.2 K x *g* in a Vortex-Genie2™ (Scientific Industries Inc. Bohemia, New York, USA) and phase separation achieved by centrifugation at 13.3K x *g* for 10 minutes. The remaining steps were followed as outlined in [44]. DNA plus a negative extraction control (nuclease-free water Qiagen, Venlo, The Netherlands) were sent to the Research Technology Support Facility at Michigan State University (Michigan, USA), for amplicon sequencing of the 16S rRNA gene targeting the V4 hypervariable region using the universal primer set (515f/806r; [45]) with indexed fusion primers to generate amplicons compatible for multiplexed Illumina sequencing. PCR products were normalised using an Invitrogen SequalPrep DNA normalisation plate and the normalised products per individual sample were then pooled to create an equimolar 16S V4 library. This library was quality checked using the Qubit dsDNA assay (Life Technologies, Darmstadt, Germany), Caliper LabChipGX (Caliper Life Sciences, Inc. Waltham, Massachusetts, USA) and Kapa Biosystems qPCR assay (Sigma-Aldrich, St. Louis, Missouri, USA). The library was then sequenced by Illumina Miseq at Michigan State University (Michigan, USA) using a standard flow cell and 500 cycle v2 reagent cartridge (Illumina Inc., Hayward, California, USA).

##### Sequencing Analysis

Raw sequences were submitted to the SRA database under Bioproject Submission PRJNA636649. A total number of 12,655,273 reads were obtained from 68 samples. The subsequent paired-end reads were demultiplexed and converted to FastQ format sequence files for further analysis outlined below. The sequence reads were filtered using Sickle (v1.2; [46]) by applying a sliding window approach and trimming regions where the average base quality drops below 20. Pandaseq (v 2.4; [47]) was used with a minimum overlap of 20 bp to assemble the forward and reverse reads into a single sequence spanning the entire V4 region [47]. After obtaining the consensus sequences from each sample, UPARSE (v7.0.1001; [48]) was used for operational taxonomic unit (OTU) construction. The sequencing analysis protocol can be accessed as outlined in the data availability section. The approach was as follows: reads from different samples were pooled together and barcodes were added to keep an account of sample origin. Reads were then de-replicated (output - 3,264,891 reads) and sorted by decreasing abundance and singletons were discarded (output - 411,123 reads). Reads were then clustered based on 97% similarity, which followed by *de novo* chimera removal from most abundant sequences (output - 3,894 reads). Additionally, a reference-based chimera filtering step using a gold database [49] was employed to remove chimeras that may have been missed in the previous step (output - 3,612 reads). This left 3,612 clean operational taxonomic units (OTUs). The assign_taxonomy.py script from the Qiime workflow [50] was used to taxonomically classify the representative OTUs against the SILVA SSU Ref NR database release (v123; [51]). These taxonomic assignments were then integrated with the abundance table using the make_otu_table.py function from the Qiime workflow to produce a biom file. To find the phylogenetic distances between OTUs, the OTUs were multisequence aligned against each other using mafft [52] and then FastTree (v2.1.7; [53]) was used on these alignments to generate an approximate maximum-likelihood phylogenetic tree in NEWICK format.

### Data Analyses

#### Calculation of Fish Condition

Fulton’s Condition Factor (K) growth parameter was calculated from the equations by recording fish length (cm) and fish body weight (g). This was carried out for a) tank populations, b) gut morphology sampled fish and c) gut microbiota sampled fish.

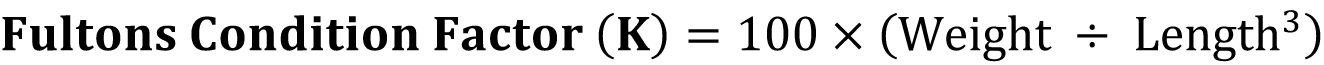

#### Microscopy Analysis

ImageJ software (version 1.46, [54] was used to measure the following light microscopy gut parameters (µm): (1) mucosal-fold height (measured from the tip to the base of the mucosal fold) and (2) lamina propria width. The microvilli (MV) density measurements taken using SEM were processed by initially cropping images to a standard size and transforming to the 8-bit image. Using the threshold function, an arbitrary value (arbitrary units, AU) for the ratio of white (MV) to black (spaces between the MV) was calculated for each image [42].

#### Statistical Analysis for Growth and Histology

Statistical analysis on the experimental trial growth data for tank populations was carried out using one-way ANOVA with a posthoc Tukey test to discern statistical differences. For the histology data (mucosal fold length, microvilli density and lamina propria width) was compared using the non-parametric Kruskal-Wallis test within Prism software (v6.03, GraphPad, San Diego, California, USA). This compared the mean rank of each treatment with the mean rank of every other treatment. A significance value (*P*) of 0.05 or under was used unless otherwise stated. This methodology was also used to determine the statistical differences in sampled fish condition (K), fish weight (g) and fish length (cm) for both gut morphology and gut microbiota sampled fish (tested separately – Figure S2).

#### Microbiome 16S rRNA Sequencing Statistical Analysis

The microbial community of the hindgut of samples across treatments was characterised to assess whether treatment impacted the microbial composition of the gut. Samples were grouped into defining categories bases on treatment and/or time (Week 0, CTRL_8, ULVA_8, ASCO_N_8, ASCO_LG_8, CTRL_12, ULVA_12, ASCO_N_12, ASCO_LG_12). These groupings were then used to investigate changes in the microbiota with respect to treatment and time. Microbiome statistical analyses were carried out via the software R (v 3.4.2[55]) with the OTU biom file, phylogenetic tree and the associated metadata files for the study. Spurious OTUs and OTUs found in the negative control were removed from all samples before statistical analyses.

##### Alpha and Beta-diversity Statistical Analyses

were carried out using R’s ‘Vegan’ package [56]. The following alpha diversity measures were then calculated on rarefied microbiome data - Pielou’s evenness (how equal in number the relative OTUs are); Richness (the number of unique OTUs) and Shannon’s index of entropy (the degree of uncertainty within a grouping assuming that a high degree of uncertainty corresponds to a highly diverse sample [57]. Pair-wise ANOVA *P*-values were drawn on top of alpha diversity figures. Beta-diversity measures were visualised using non-metric multidimensional scaling (NMDS) plots using standard dissimilarity distance measures: Bray-Curtis and Unifrac (Unweighted and Weighted). Unifrac distances were calculated using the R package ‘Phyloseq’ [58]. Samples were grouped for different treatments as well as the mean ordination value and spread of points (ellipses were drawn using Vegan’s ordiellipse() function that represents the 95% confidence interval of the standard errors). R’s ‘Vegan’ package and ordisurf() function was used to determine if the main pattern in species composition described in principal co-ordinate analysis (PCOA) could be explained by the variable of fish condition factor (K). Ordisurf() internally fits a generalised additive model “*gam*” smoothing with formula K ~ s(PCoA1, PCoA2).

##### Characterising the microbial profiles across treatment groups and over time

In order to interpret this complex high-throughput dataset we used R’s mixOmics package [59] and Sparse Projection to Latent Structure – Discriminant Analysis (sPLS-DA) to select the most important and distinguishing OTUs between the treatments and over time. Analysis was performed at genus level with genera > 3.5% abundance and with TSS + CLR (Total Sum Scaling followed by Centralised Log Ratio) normalisation applied. OTU feature selection and multiple data integration was achieved by constructing artificial latent components (inferred variables) of the OTU table by first factorizing the abundance table and response variable (that contains categorical information of samples) matrices into scores and loading vectors in a new space such that the covariance between the scores of these matrices are maximised. In doing so, the algorithm also enforces a constraint on the loading vector associated with the abundance table (that has corresponding weight for each feature) such that some components of the loading vector go to zero and only significant features in terms of their discriminatory power remain. In this study, sPLS-DA was applied to determine the effect of ‘*Treatment*’ and ‘*Time*’. ‘*Time*’ contained Week 0 and all treatments except the ASCO_LG fish (i.e. CTRL, ULVA, ASCO_N) and longitudinal information (Week 0, Week 8, Week 12). ‘*Treatment*’ examines the independent treatments including all groups (CTRL, ULVA, ASCO_N and ASCO_LG) at either Week 8 or Week 12. The number of latent components and the number of discriminants were calculated. In finding the number of discriminants the model was fine-tuned using leave-one-out cross-validation and subsequent balanced error rates which accounts for sample number differences (fine-tuning parameters indicated in figure legends for Figure 6 and Figure S5). The final output was a heatmap containing the differential genera over the number of tuned components which is discussed in the Results section. Further description of the implementation of sPLS-DA for microbial data-sets can be found in Gauchotte-Lindsay et al. [60]. The OTUs identified in sPLS-DA were subsequently used in differential tree analysis to show the taxonomic shifts in microbial community structure occurring over time. The complete methodology is outlined in [61] and uses the DESeqDataSetFromMatric() function from the DESeq2 package [62] Metacoder package to generate the tree visualisations [63] in R.

##### Finding a core microbiome and core microbiome per treatment

We followed a method for identifying the core microbiome as outlined McKenna et al. [60], and Shetty et al. [63] using R’s microbiome package [65] adjustable parameters for percentage relative abundance and percentage prevalence within samples. For the samples in this study, the core microbiome was defined as prevalent in 85% of samples [66].

## Results

### Fish Growth, Condition and Survival

Mean weight of Atlantic cod juveniles increased from 123g at the start of the feeding trial to 236 ±62g, 243 ±72g and 190 ±84g (SD) for the CTRL, ULVA and ASCO treatment total tank populations, respectively at the end of the trial. An almost two-fold increase in weight was measured in the fish fed the ULVA and CTRL diets. The analysis showed there were significant (*P < 0.05*, F = 73.53, df=2) differences between the dietary groups, with a Tukey test revealing ASCO fish weighed less than CTRL and ULVA groups. *Note:* The ASCO treatment was not split into ASCO_N and ASCO_LG for tank population data. All tank populations had individual fish with condition factor of <0.85. The prevalence of fish with condition factor <0.85 in the ASCO treatment was 33%, as compared to just 3% in the CTRL and 8% in the ULVA treatment at Week 12. Correspondingly, the ASCO treatment had a greater percentage mortality rate at 21%, while the CTRL and ULVA treatments had values of 19 and 16%, respectively.

Within the fish sampled for microbiology and histology, statistically significant differences (Kruskal-Wallis; *P* < 0.05) were observed in total weight (g) and condition factor (K) of the ASCO_LG group as compared to the remaining treatments (Supplementary Figure S2a and S2b). At Week 8 within the microbiology samples, the ASCO_LG group had a condition factor of 0.77 ±0.02 (SD), compared to 1.16 ± 0.14 (SD) in the ASCO_N group. In addition, there were no significant differences (*P* > 0.05) found between the weight or condition factors for the CTRL, ULVA or ASCO_N treatments (229-256 g fish-1 and condition factor of 1.16-1.21). This trend continued in Week 12 with a further decrease in condition factor in the ASCO_LG group observed.

### Intestinal morphology

In the ULVA and CTRL groups, the microvilli were in a densely packed arrangement of uniform shape (Figure 1). In contrast, both ASCO groupings (Normal and Lower Growth) had a more heterogeneous structural arrangement. Analysis of Week 8 data indicated that the hindgut of the ASCO_LG grouping had significantly reduced mucosal fold height (116 ±42 µm, *P <0.0005*) and significantly increased lamina propria width (106 ±44 µm *P <0.01*) compared to 220 ±60 µm and 69 ±28 µm in the CTRL for height and width, respectively (Table 2). It was noted upon dissection even after the starvation period that undigested material was contained in the digestive tract of fish within this group. There was no significant difference between the lamina propria width and mucosal fold height of the ULVA group compared to the CTRL group (*P >0.05* and *P >0.05*, respectively). A similar trend was observed at Week 12, with ULVA and CTRL groups not being significantly different in terms of lamina propria width and mucosal fold height. Interestingly, the ULVA diet had increased microvilli density (AU) when compared to the CTRL, ASCO_N and ASCO_LG groupings (Figure 1 and Table 2). However, this relationship was only significant between the ASCO_N at Week 8 (*P* < 0.01) and ASCO_LG at both Week 8 (*P* < 0.01) and Week 12 (*P* < 0.05).

**Table 2.** Gut morphology from hindgut sections taken from juvenile Atlantic cod (*Gadus morhua*) from the CTRL, ULVA, ASCO_N and ASCO_LG groupings at Week 8 and Week 12. Mucosal-fold length (µm) and laminar propria width (µm) measurements are taken from light microscopy images while the microvilli density (AU) measurements are taking from scanning electron microscopy images (± SD). The associated statistical measurements indicated are calculated from Kruskal-Wallis significance testing *P* < 0.05*, *P* < 0.01**, *P* < 0.001***.

|  | Week 8 |  |  | Week 12 |  |  |
| --- | --- | --- | --- | --- | --- | --- |
|  | Mucosal-fold length | Laminar propria width | Microvilli density | Mucosal-fold length | Laminar propria width | Microvilli density |
| CTRL | 221.06 (± 32.04) | 58.58 (± 12.04) | 4.07 (± 0.66) | 152.48 (± 34.92) | 31.43 (± 4.85) | 4.23 (± 0.52) |
| ULVA | 178.48 (± 19.56) | 70.14 (± 20.65) | 4.89 (± 0.44) | 164.11 (± 23.57) | 33.35 (± 5.50) | 4.85 (± 0.81) |
| ASCO_N | 152.43 (± 38.89) | 102.17 (± 22.46) | 3.67 (± 0.30) | 187.42 (± 39.28) | 38.26 (± 9.11) | 4.02 (± 0.72) |
| ASCO_LG | 115.61 (± 3.173) | 106.19 (± 18.82) | 3.47 (± 0.53) | 95.62 (± 16.16) | 48.75 (± 9.91) | 3.64 (± 0.42) |
| Statistics | *CTRL+ASCO_N<br>***CTRL+ASCO_LG<br>*ULVA+ASCO_LG | *CTRL+ASCO_N<br>**CTRL+ASCO_LG<br>*ULVA+ASCO_LG | ** ULVA+ASCO_N<br>**ULVA+ASCO_LG | *CTRL+ASCO_LG<br>*ULVA+ASCO_LG<br>***ASCO_N+ASCO_LG | **CTRL+ASCO_LG<br>*ULVA+ASCO_LG | *ULVA+ASCO_LG |

**Figure 1.**
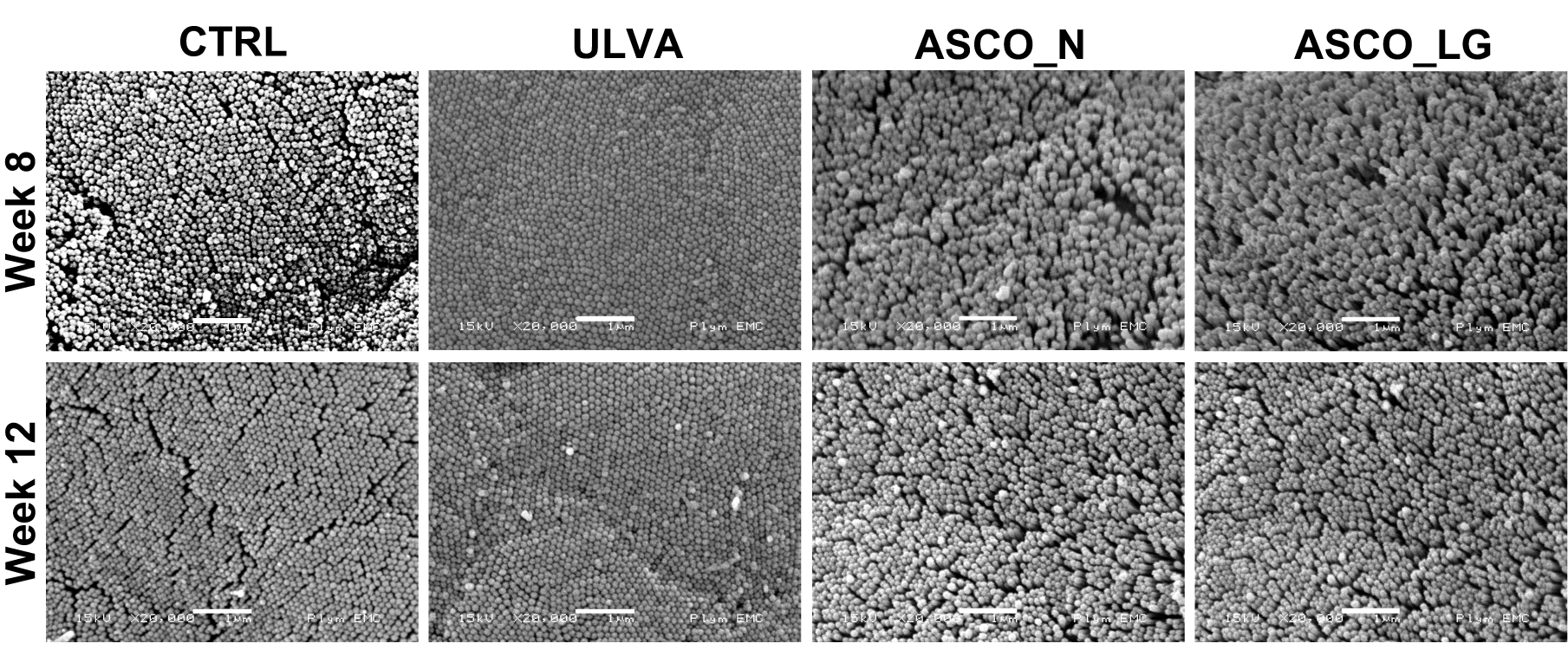
Scanning electron microscopy images of the hindgut of juvenile Atlantic cod (*Gadus morhua)* from each treatment (CTRL, ULVA, ASCO) at Week 8 and Week 12. Scale bar represents 1 µm.

### Hindgut Microbial Community Composition

The hindgut of Atlantic cod (*Gadus morhua*) was characterised using amplicon sequencing of the 16S rRNA V4 region. The data presented here are based on the sequences from 67 samples obtained following quality filtering as described in the Methods section. A total of 12,655,273 reads were generated, resulting in 411,123 quality-filtered paired-end reads once singletons were removed, corresponded to 3,612 unique OTUs at the 97% similarity level. The taxonomically assigned data (> 97% similarity) showed that members of the *Proteobacteria* and *Firmicute*s phyla were dominant across all samples comprising as much as 98% and 90% relative abundance respectively (Figure 2A). The phylum *Bacteroidetes* increased over time to as much as 80% relative abundance in the CTRL fish at Week 12. The most frequently observed OTUs (Figure 2B) were OTU_307 and OTU_372, both *Photobacterium* spp. Over time OTUs identified as Bacteroidetes increased - OTU_9 [*Bacteroides* sp.] and OTU_20 [*Alistipes* sp.] appeared in the Top 25 phyla. The ASCO_LG samples at Week 12 (ASCO_LG 12) showed a high abundance of OTU_1968 [*Psychromonas* sp.]. A large proportion of the OTU-level taxonomically assigned data was assigned to the category ‘Others’, i.e. less abundant genera, as much as 99% in some Week 0 samples (Figure 2B).

**Figure 2.**
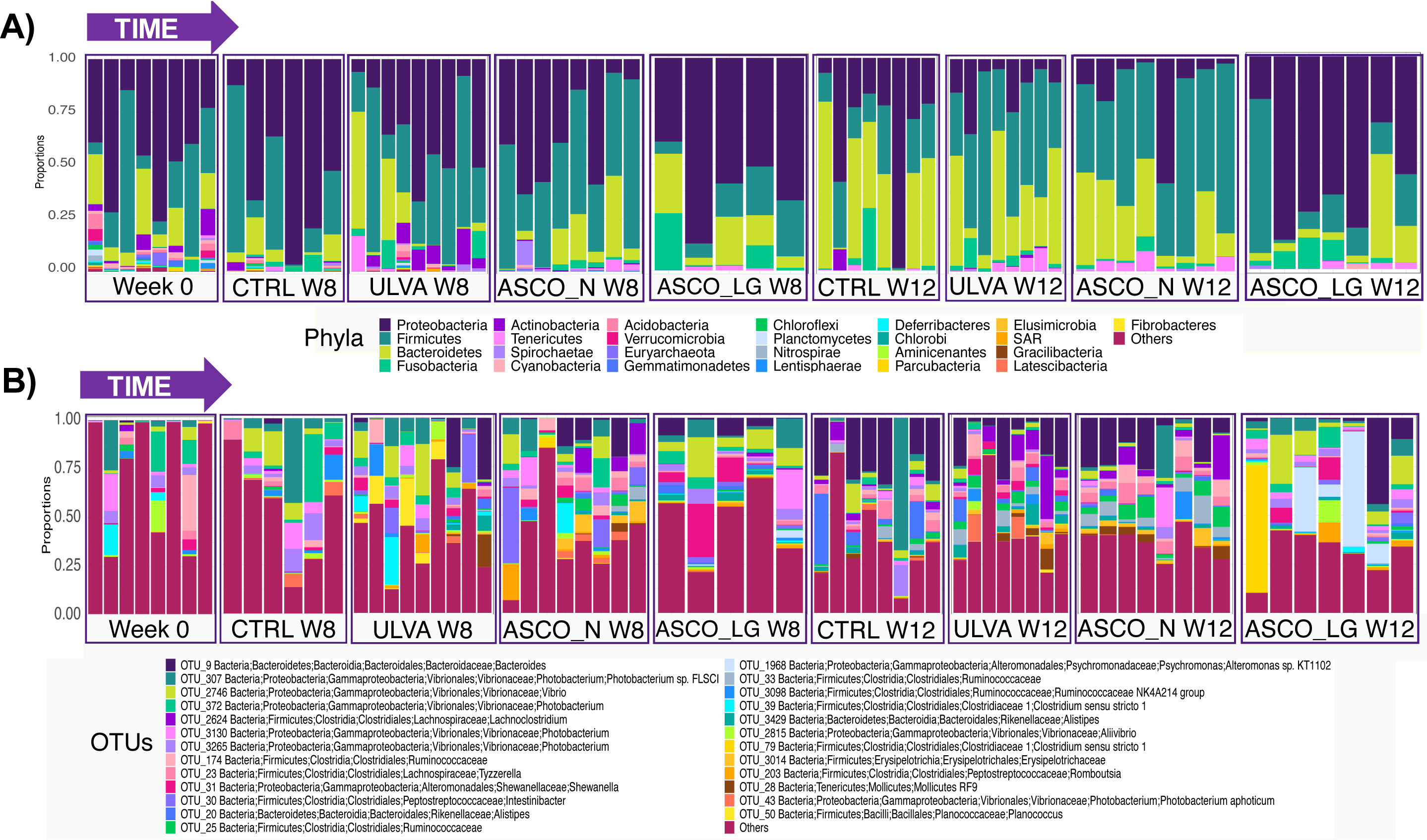
Horizontal bar-plot showing the relative abundance of the top 25 most abundant, **A**) phyla, and **B**) Operational taxonomic units (OTUs) in the hindgut of juvenile Atlantic cod ordered by the dietary groups over time. “Others” represents the remaining lesser abundant taxa.

### Diversity of the Hindgut Microbial Communities

The alpha-diversity measures used in this study included the following a) species evenness - how similar in numbers each species is, b) species richness - the number of different species, and c) Shannon index - a measure of entropy *i.e.* the degree of uncertainty as a proxy for diversity. A significant decrease in all these tested measures was observed from Week 0 to Week 12, irrespective of dietary treatment (*P* <0.05, Figure 3A and Supplementary Information Figure S3). No significant difference was observed between Week 8 and Week 12 samples (excluding ASCO_LG samples). The ASCO_LG fish at Week 12 were less rich than those from the CTRL (*P* <0.05), ULVA (*P* <0.05) and ASCO_N (*P* <0.001) individuals from the same time point (Figure 3A).

**Figure 3.**
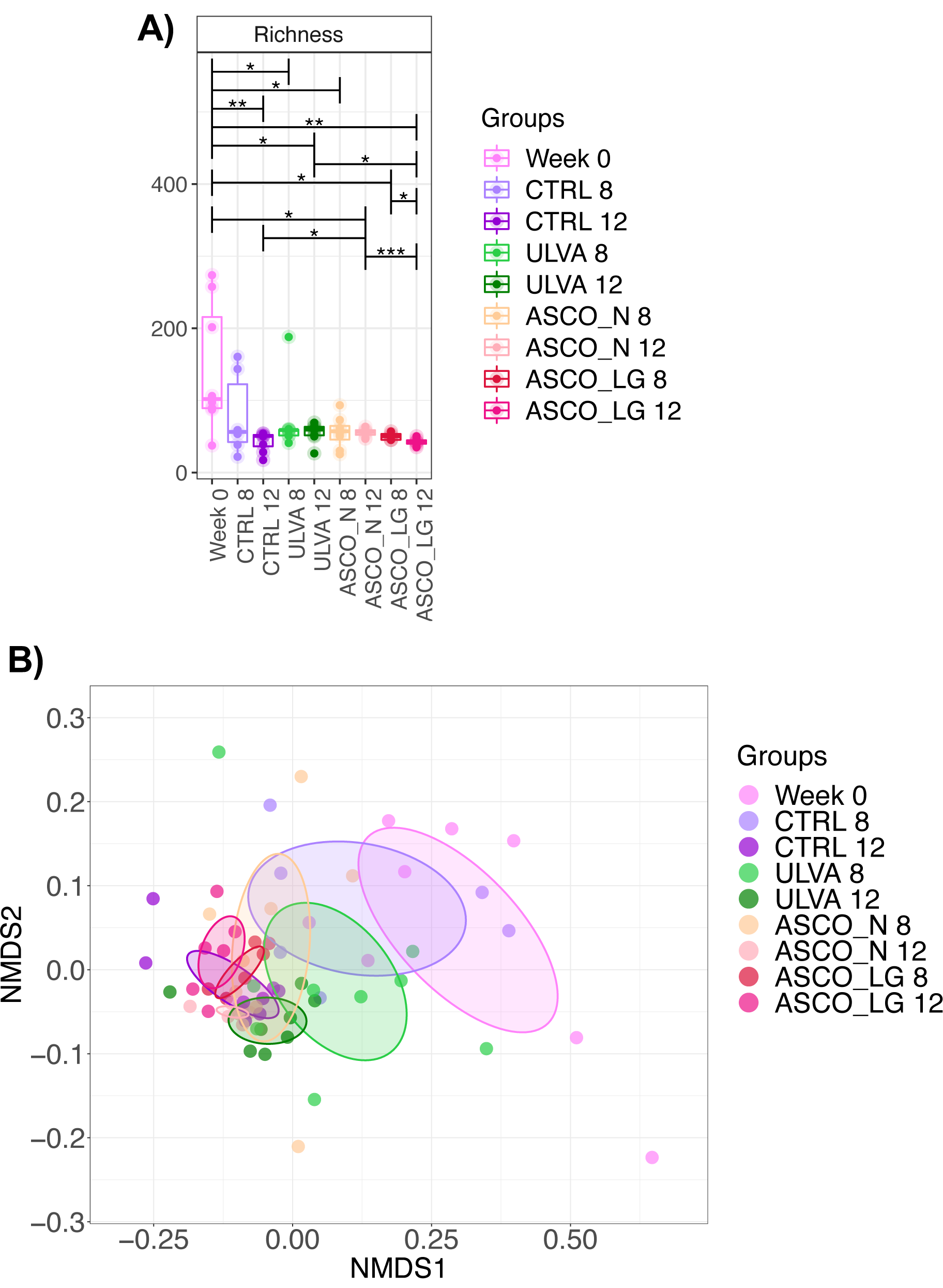
**A)** Alpha diversity measure: Species Richness with the corresponding colour legend for the treatment groups. Pair-wise ANOVA *P*-values are displayed *P* < 0.05*, *P* < 0.01**, *P* < 0.001***. **B)** Non-metric distance scaling (NMDS) plot based on Unifrac phylogenetic distance with colour coded circles representing the differing treatment groups over time (R^2^ = 0.27182, *P*= 0.001).

We observed a clear differentiation composition of the microbial community in the beta-diversity measures between Week 0 samples and the CTRL and macroalgae dietary supplement groups at Week 8 and Week 12 (Figure 3B). The microbial communities across all three treatments (CTRL, ULVA, ASCO_N, ASCO_LG) became increasingly similar over time – forming overlapping clusters (irrespective of treatment) by Week 12 (Figure 3B). Dissimilarity observed could partly be explained by variation in OTUs of low abundance as weighted Unifrac (phylogenetic distance weighted by abundance counts) analysis showed all treatments and time points within overlapping clusters (Supplementary Information Figure S3). Despite showing a pattern of convergence over time treatment groups (Week 0, CTRL, ULVA, ASCO_N, ASCO_LG at Week 8 and Week 12) were significantly different in all beta-diversity non-metric multi-dimensional scaling plots (*P* < 0.001).

### Temporal changes in the hindgut microbial community

The data above indicates that the hindgut microbial communities were strongly influenced by time, as opposed to treatment. Over time, the number of unique OTUs reduced from 151 at Week 0 to only 12 that were unique at Week 12 (Supplementary Information Figure S4). To identify what OTUs were changing over time ‘differential’ analysis was used. Discriminating OTUs between Week 0 and treatments (excluding ASCO_LG) at Week 8 and Week 12 (CTRL, ULVA and ASCO_N) were identified using sPLS-DA and displayed in a heatmap (Supplementary Information S5). The microbial profile of Week 0 fish was distinct from the subsequent time points and the discriminating OTUs from Week 0, Week 8 and Week 12 were used to create differential trees to visually demonstrate the taxonomic changes over time (Figure 4). The differential OTUs at Week 0 and Week 8 contain many *Xanthomonadales*, *Rhodobacteriales*, *Desulfovibrionales*, *Mollicutes*, *Gemmatimonodales*, *Clostridiaceae*, and *Bacilli* taxonomic nodes. While Week 12 taxonomic nodes consisted of *Ruminococcaceae*, *Lachnospiraceae*, *Christensenellaceae*, *Erysipelotrichia*, and *Bacteroidetes* members.

**Figure 4.**
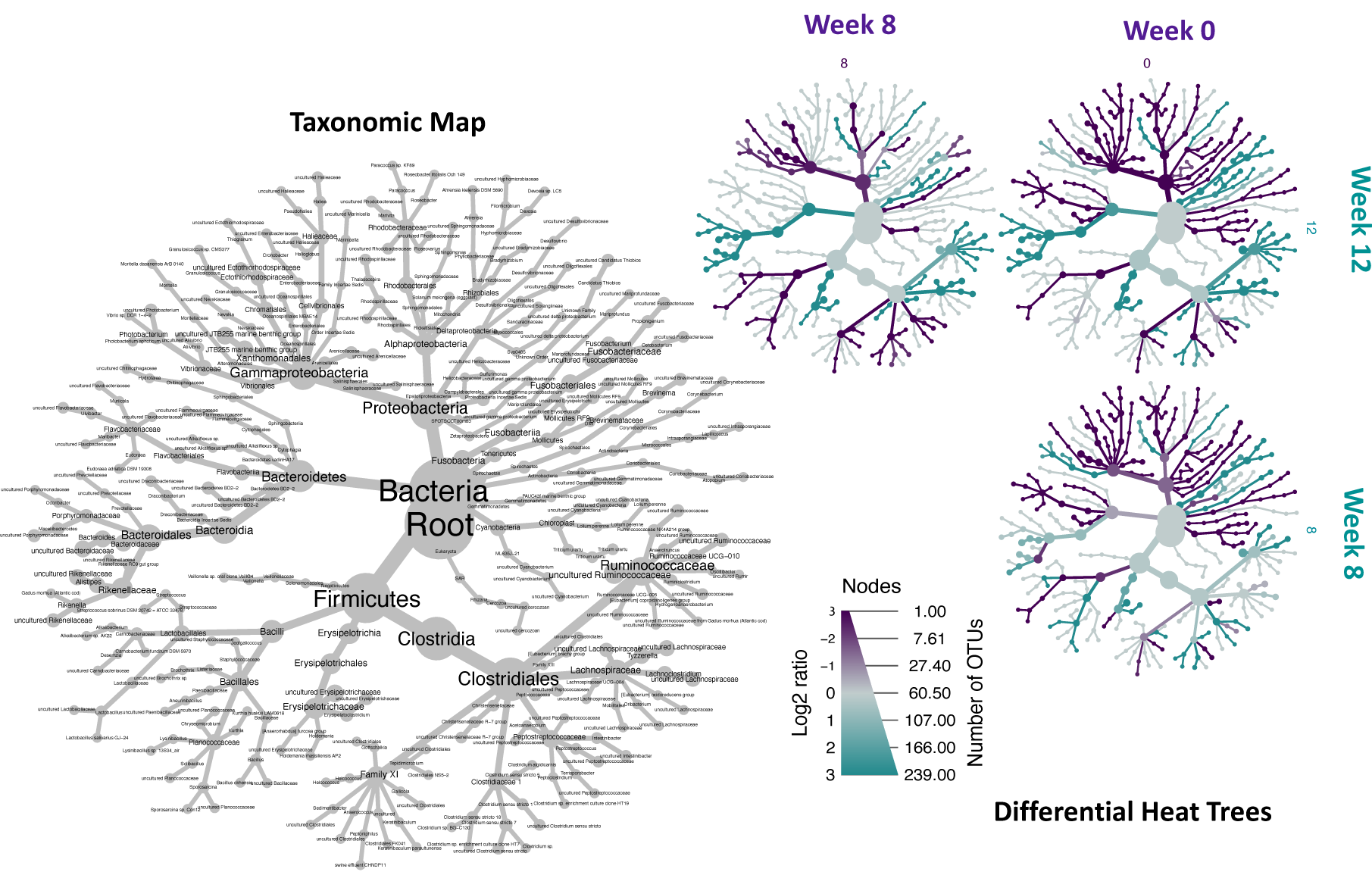
Differential taxa in the hindgut microbiota of juvenile Atlantic cod (*Gadus morhua*). These taxa were first determined using sPLS-DA with the discriminating operational taxonomic units (OTUs) at > 1% relative abundance at Week 0, Week 8 and Week 12 (excluding ASCO_LG fish). The taxonomic map is a legend to show the identity of the differentiating taxa. This legend is read to inform the differential heat trees. The differential heat trees show the microbiota shifts from Week 0, Week 8, and Week 12. The branches where they are upregulated are coloured according to their respective categories shown for each tree (Week 0, Week 8, and Week 12). The nodes legend can be read as follows: Left side ‘Log2 ratio’ of the median proportions of the taxa corresponding to the colour scale, Right side ‘Number of OTUs’ where the diameter reflects the number of OTUs corresponding to the nodes on the heat trees.

### Influence of macroalgae supplement on the hindgut microbiome

#### Changes in diversity across treatments

The microbial diversity in the hindgut of juvenile cod decreased over time and the microbial communities became increasingly similar over time. Physiologically, however, significant changes were occurring with respect to the fish condition and the ASCO group were categorised into ASCO_N and ASCO_LG groupings from Week 8. Differences in beta-diversity among the groups was tested at Week 8 and Week 12 (Local Contribution to Beta-Diversity measurement), with no significant difference found across all dissimilarity matrices. This was with the exception of the ASCO_N samples which were significantly different to all treatments using Unifrac phylogenetic distance at Week 12 (CTRL; *P* = 0.019, ULVA; *P* = 0.47, ASCO_LG; *P* = 0.002). To understand if the fish condition could be linked to microbial community composition patterns Unifrac phylogenetic distance of OTUs was used and compared using Principal Coordinates Analysis (PCoA) plots containing contour maps with the condition factor values (Figure 5). The culminative explained variance was as follows; Week 8 - Dim1 = 20.55% and Dim2 = 8.77%, Week 12 - Dim1= 17.9% and Dim2 8.81%). At Week 8, ASCO_LG fish clustered around contour values of ~1. A clear distinction was not observed between treatment groups at this time point. ASCO_LG fish condition decreased from Week 8 to Week 12, and this corresponded with a response in the microbial community structure as evidenced by ASCO_LG individuals at Week 12 clustered around condition factor contours of values <0.75 while the remaining groups (CTRL, ULVA and ASCO_N) clustered together with values generally above 1.

**Figure 5.**
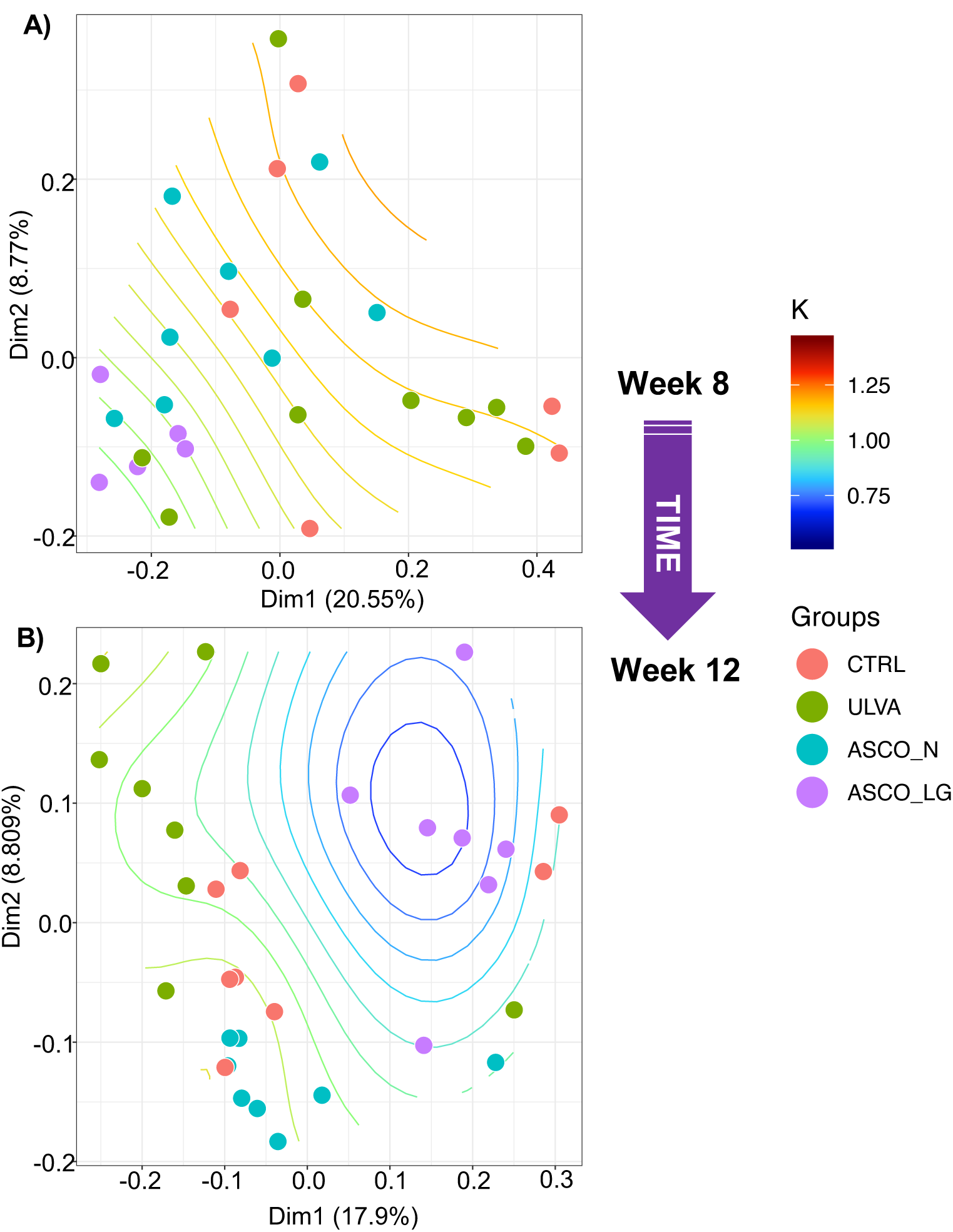
Unifrac unweighted principal co-ordinate analysis of the gut microbiota of juvenile Atlantic cod (*Gadus morhua*) community composition based on the different dietary treatments (CTRL, ULVA, ASCO_N and ASCO_LG) at **A**) Week 8, and **B**) Week 12. The values given on each axis (Dim1 or Dim2) indicate the total percentage variation for the treatment groups. Contour lines represent condition factor (K) fit through generalised additive modelling. Deviance in ordination space at Week 8 is explained by 24.5%; R^2^ = 0.18; * *P* = 0.043. Deviance in ordination space at Week 12 is explained by 57.8%; R2 = 0.5; *** *P* = 0.0002.

#### Differential OTUs across treatments

sPLS-DA was used to discriminate the specific OTUs which were influenced by the dietary treatments and therefore determine the influence of macroalgae supplement on the hindgut microbiome. This was performed with the Week 0 samples removed and Weeks 8 and 12 were assessed separately to exclude any artefacts with respect to temporal sampling. The macroalgal treatments were compared to the corresponding CTRL group and the information displayed in a heatmap. Week 8 sPLS-DA comparison is contained within Supplementary Information Figure S6. At Week 12, the ASCO_LG group formed a separate branch (Figure 6), while the remaining treatments (CTRL, ULVA, ASCO_N) clustered together. The differential OTUs between treatments formed three large phylogenetic branches. The bottom branch contained OTUs (*Oscillibacter*, *Erysipelotrichaceae* and *Rikenellaceae* spp.) primarily downregulated in the ASCO_LG fish compared to the other diet treatments. The middle branch contained OTUs (*Psychrosomonas* spp., *Propionigenium*, *Clostridium sensu stricto*, and *Rhodospirillaceae*) primarily upregulated in the ASCO_LG fish compared to the other diets. The microbial profiles of the CTRL, ULVA, and ASCO_N groups showed a high degree of similarity within this plot.

**Figure 6.**
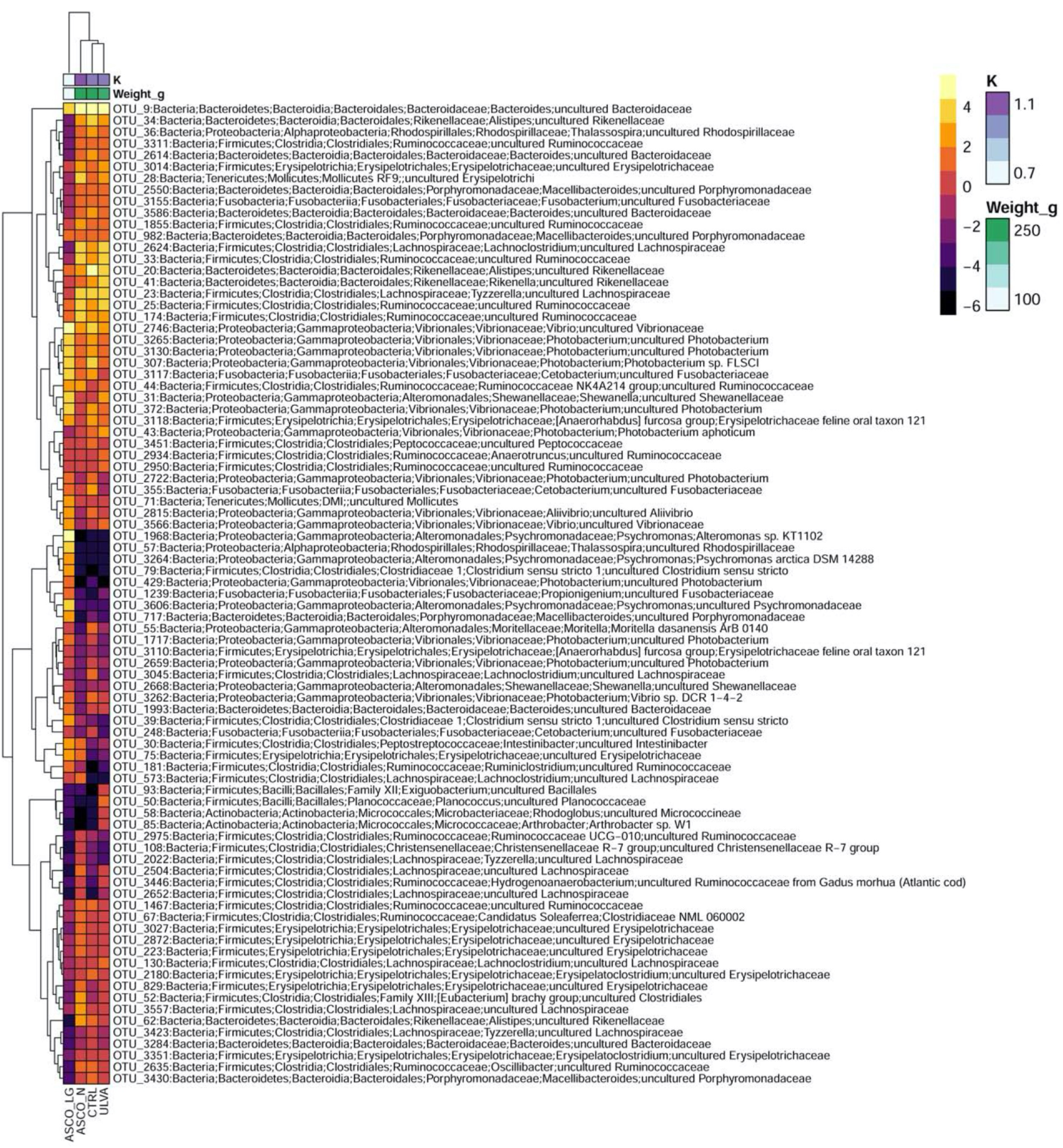
sPLS-DA heatmap of the discriminant operational taxonomic units (OTUs) > 3.5% relative abundance between the CRTL, ULVA, ASCO_N and ASCO_LG groups of juvenile Atlantic cod (*Gadus morhua*) at Week 12. Rows and columns are ordered using hierarchical (average linkage) clustering to identify blocks of genera of interest. The heatmap depicts TSS+CLR (Total Sum Scaling followed by Centralised Log Ratio) normalised abundances: high abundance (orange/yellow) and low abundance (dark purple). Fish weight (g) and condition factor (K) is shown per grouping. sPLS-DA was fine-tuned using centroids.dist and 3 tuning components.

#### Core Microbiome of Control diet versus Ulva rigida supplement

The data highlighted above shows that the differences over time were a stronger influence than treatment. The core microbiome (found in 85% of individuals) across diets and treatments consisted of just 3 OTUs – OTU 2746 [*Vibrio* sp.], OTU 307 and 2746 [*Photobacterium* spp.]. Given the palatability issues observed with the *A. nodosum* supplement, it is unlikely that this would be used in its current form as a feed supplement. As such, in the following section, the core microbiome was compared for the CTRL and ULVA fish only at Week 12 (Supplementary Information Figure S7). The core microbiome of the CTRL group consisted of 8 OTUs listed in order of abundance and prevalence (OTU 9 [*Bacteroidetes* sp.], OTU 2746 [*Vibrio*], OTU 307 [*Photobacterium* sp.], OTU 2624 [*Lachnoclostridium* sp.], OTU 3265 and 3130 [*Photobacterium* spp.] and OTU 982 and OTU 2550 [*Macellibacteroides* spp.]). The fish fed on the ULVA diet shared all OTUs within the CTRL group, except for OTU 3265 and OTU 3130, which were identified as *Photobacterium* spp. In addition, the ULVA group contained OTUs not found in the CTRL group. These were broadly identified as *Ruminococcaceae* spp., *Shewanella*, *Tyzerella, Rikenellaceae* spp., *Peptococcaceae*, *Bacteroides*, and *Fusobacterium*.

## Discussion

In this work we evaluated the feasibility of supplementing juvenile Atlantic cod (*G. morhua*) diets with 10% macroalgae with specific consideration to the health of the intestine. To the author's knowledge, few publications have considered the impact of macroalgal dietary supplementation on the gut physiology and microbiome in cod. Our research study aimed to address the following knowledge gap, examining two different macroalgal species: *U. rigida* and *A. nodosum* on fish condition and hindgut microbiome.

The data presented in this study indicated that a diet fed to juvenile Atlantic cod could be supplemented with 10% *U. rigida* without negative impacts on growth performance, gut physiology, or microbiome shifts as compared to the control treatment. This is in contrast, with reports from rainbow trout (*O. mykiss*) supplemented with a different *Ulva* species (*U. lactuca*) at 10% inclusion level had poorer growth compared to those fed control diets [67]. Differences in the response to specific macroalgae ingredients across fish species is unsurprising given the variation in multiple factors including water habitat type, gut morphology, digestive enzymes, and fish genetics. This highlights that extensive research is needed for individual fish species to ascertain the suitability of specific ingredients. Our positive result correlates with studies on European sea bass (*D. labrax*; [68] and Senegalese sole (*Solea senegalensis*; [69]), indicating that *U. rigida* can be supplemented in the diets of a variety of finfish species. *U. rigida* is ubiquitous and can be a nuisance species, proliferating in large quantities resulting in green tides. It is this significant algal biomass availability in the wild that would provide the upscale requirement in commercial aquafeed production [23].

Based on the current study, *A. nodosum* was determined not to be a suitable dietary supplement for juvenile Atlantic cod at 10% inclusion levels. In a subset of the population, growth, fish condition, intestinal physiology and the gut microbiome were all impacted by this dietary inclusion. Individuals that tolerated the *A. nodosum* supplement (ASCO_N) had similar condition factors to those in the CTRL and ULVA dietary groups. There appeared to be two reasons for fish not tolerating the *A. nodosum* supplement: palatability and digestibility. The ASCO diet was unpalatable to many fish within this treatment, evidenced by fish rejecting pellets (‘spit outs’) during a feeding event. Palatability issues has been documented in many fish species, including Atlantic cod fed a microalgal supplement [70]. Taste experiments with the common periwinkle have shown a preference for *Ulva* species over *A. nodosum*, even during starvation [71]. Cod is a benthic species, living close to the seafloor, and uses an enhanced chemoreceptor system with a large number of external cutaneous taste buds on areas like the pelvic fin to locate prey [72][73]. Thus, they may have heightened sensitivity to stimuli or antinutritional factors (ANF). Moreover, ANFs (e.g. phenolics, polysaccharides, and tannins) in the supplement ingredients can have a bitter taste [74]. These ANFs are also known to influence substrate digestibility [75]. During dissection, it was noted that the fish in the ASCO treatment had undigested material at the rear of the digestive tract. This was particularly prominent in the ASCO_LG fish. The ASCO_LG fish not surprisingly had significantly reduced weight and condition factor compared to the ULVA, CTRL and ASCO_N groups. These results correlate with work by Yone et al. [82] where 10% *A. nodosum* supplementation decreased the final weight and feed efficiency in red sea bream.

Gut morphology plays a key role in the digestive function and overall health of a fish species. Absorption of nutrients is mediated by the function and morphology of the gut. The microvilli sit on a single layer of epithelial cells (enterocytes) which facilitates the transport of nutrients through the gut wall barrier. Healthy microvilli (long and densely packed) are proposed to be associated with improved nutrient uptake due to increased surface area for absorption [78], [79]. It is well established that alternative aquafeed ingredients can impact intestinal morphology [80]. Thus, changes in the cod hindgut microvilli density or mucosal-fold length in response to a change in diet could reveal potential changes in nutrient absorption. In this study, the microvilli density, mucosal fold height and lamina propria width in the ULVA group had higher mean values than all groupings (CTRL, ASCO_N and ASCO_LG), though a significant difference was only observed in the case of the ASCO fish. A similar finding was reported by Moutinho et al. [69] for Senegalese sole (*S. senegalensis*) showing a trend towards higher mucosal fold height but this was not significant when compared to the control. The ASCO_LG fish had an altered hindgut morphology. These alterations included, fatter laminar propria, decreased mucosal-fold height and a modest reduction in microvilli density. Thickening of the lamina propria is suggested to be due to the infiltration of inflammatory cells [81]. This may have increased the susceptibility of these fish to potential disease and predation by larger juveniles [82]. Though we observed no clinical signs of disease in any of the dietary groups.

In some aquafeed studies, supplementing the diet with plant ingredients (soybean, pea and canola) has resulted in negative impacts on fish gut morphology and reduced growth with combined disruption to the gut microbiota as observed in rainbow trout [83]. While in contrast, in a replacer study (where the complete protein source is replaced) no change in the gut microbiota was observed despite substantial changes in gut morphology (e.g. Atlantic salmon soybean replacer study [84]). Recent research on a variety of fish species has indicated that gut microbiota can be linked to digestion [30], immunity [85], [86], and disease prevention. Therefore, this is an important parameter to consider when determining the impacts of diet supplementation. We showed that macroalgae supplementation did not impact the hindgut microbiome. Rather, time was a greater driver with hindgut microbial communities converging to be more similar over time irrespective of treatment. Indeed, before the start of the experiment, the fish were fed the same diet (commercial fish pellet) and showed the greatest difference in community composition among individuals. Over the course of the experiment, as the juvenile cod grew, the microbial diversity in all treatments decreased and despite differences in the feed the microbial community structure converged among treatments over the 12-week period. These changes were accompanied by the differential increase and simultaneous decrease of a set of key OTUs (25 OTUs) over the course of the experiment. In fact, over the 12-week period the number of unique OTUs decreased from 152 at Week 0 to just 12 at Week 12. In terms of composition, the early hindgut microbiome was dominated by *Proteobacteria* and *Firmicutes*. The dominance of these phyla is not surprising as these bacteria are ubiquitous. In the context of the fish gut microbiome, Proteobacteria and Firmicutes have been found in high abundances in the gut of many omnivorous and carnivorous fish species. These have ranged from diverse environments [87] and coastal seawater samples [88]. The reason for their dominance has yet to be fully understood. However, over time, the relative abundance of *Bacteroidetes* members increased across all treatments, except for the ASCO_LG fish group. By week 12, Bacteroidetes was the dominant phyla present in the hindgut of the majority of fish. Within gut environments *Bacteroidetes* can outcompete other phyla due to their metabolic flexibility and tractability in response to host pressures, e.g. the host immune system and gut environment pressures such as low pH [89]. Interestingly our work contrasts with the finding by Xia et al. (2014) where 12 days of starvation led to a shift towards *Bacteroidete* members in the gut microbiome of Asian seabass (*Lates calcarifer*) [32].

At week 0 and 8, we identified the discriminating taxa as *Xanthomonadales*, *Rhodobacterales*, *Desulfovibrionales*, *Mollicutes*, *Gemmatimonadetes*, *Clostridiaceae*, and *Bacilli*, which were overrepresented in terms of differential abundance within the hindgut microbial community. While, the presence of *Xanthomonadales* are less frequently found in the fish gut microbiota, they have been identified as bacterial symbionts [90] aiding the breakdown of complex hydrocarbon sources [91]. The group *Rhodobacterales* were postulated as a key bacterial species responsible for sea cucumber growth linked to polyhydroxybutyrate (energy storage molecules) metabolism [92]. *Desulfovibrionales* species are sulfate-reducers and produce hydrogen sulfide. They are commonly found in anoxic environments, their exact role and impact on fish health is unknown but within the human gut, they are linked with gut inflammation in disorders such as ulcerative colitis [93]. *Mollicutes* are an interesting group of bacteria, lacking a cell wall, and include *Mycoplasmas* which are well-known fish parasites [94]. Some studies have proposed that *Mycoplasmas* are commensal gut or even intra-host dependent microorganisms in Atlantic salmon [29], [95]. *Gemmatimonadetes* are largely uncharacterised though have been found previously in the gut of sea bream (*Sparus aurata*) and sea bass (*Dicentrarchus labrax*), at low abundances [96]. *Clostridiaceae* (*Clostridium sensu stricto*) are common gut bacteria and are metabolically flexible, capable of metabolising a range of carbohydrates to produce alcohols and acids [97]. *Bacilli* are also common fish gut bacterial species [98] that can produce a range of hydrolytic enzymes [99]. Indeed, *Bacilli* are perceived as part of a healthy gut microbiome. *Bacilli*-based probiotics have been used as dietary supplements to fish and shellfish species [100].

By Week 12, *Ruminococcaceae*, *Lachnospiraceae*, *Christensenellaceae*, *Erysipelotrichia* and *Bacteroidetes* members were the differentially abundant species from the former time points. *Ruminococcaceae* and *Lachnospiraceae* are commonly found genera, which together are responsible for protein break down and fermentation [101]. *Christensenellaceae* species are fermentative species and common in gut microbiome studies [102]. *Erysipelotrichia* and *Bacteroidetes* are saccharolytic species [101]. These patterns tend to indicate a change in microbial functioning within the gut and interactions over time towards fermentative activity and likely not a case of functional redundancy. In contrast, the ASCO_LG fish did not follow this pattern of gut microbiome development across week 8 and week 12 had upregulated *Psychromonas*, *Propionigenium*, *Clostridium sensu stricto* and *Rhodospirillaceae* species as compared to the remaining treatments. *Psychromonas* have been proposed to provide nutritional compensation for an unbalanced diet in deep-sea leeches (*Piscicolidae*), although this link remains to be further elucidated [103]. While, *Propionigenium* is known to degrade cellulose and complex carbohydrates, and are also shown to produce anti-inflammatory products as end products that could play a beneficial role in gut health [79]. Future work to understand the ecological processes underpinning the microbial community development in juvenile Atlantic cod could shed valuable insights into the mechanisms responsible for the gut microbiota patterns seen.

It has been evidenced in European seabass (*D. labrax*) that fish across different dietary regimes have a core gut microbial community that is stable over a six-week time period. While our results indicate that a core microbial community in Atlantic cod over a 12-week period consisted of just three OTUs – identified as *Photobacterium* spp. and a general *Vibrio* sp. Interestingly, the most frequently observed OTUs in the present study were identified as *Photobacterium* spp. a gamma proteobacteria within the *Vibrionales* order, widely occurring in the marine environment. These species have been found associated with numerous fish studies [for a review see 92], and have been described as and range from proposed pathogens [106] to proposed mutualistic species for the digestion of complex polysaccharides such as chitin [30]. The ubiquitous presence of *Photobacterium* spp. as part of the core microbiome suggests it may play a role in the breakdown of food and fermentation in the hindgut of juvenile Atlantic cod. The core microbiome of the ULVA treatment at Week 12 shared the same core community. These largely consisted of members of the bacterial orders *Vibrionales* (*Vibrio* and *Photobacterium*), *Bacteroidales* (*Bacteroidetes* and *Macellibacteroides*), and *Clostridiales* (*Lachnoclostridium*). This indicates that the macroalgae supplement did not alter the core microbial community in juvenile Atlantic cod.

The overall temporal trend of converging microbial communities indicated no effect of dietary supplement (at 10% inclusion levels) on the hindgut microbiota of juvenile farmed Atlantic cod. There are no studies that consider the gut microbiome development in farmed Atlantic cod from larvae to maturation. Our work provides valuable insight into the development of the hindgut microbiome of farmed Atlantic cod in the juvenile life cycle stage. It has previously been shown in other fish, that the gut microbiota is highly malleable during the first-feeding and initial lifecycle period (e.g. rainbow trout [107]), and stabilises over time as we observed. The microbial associations observed need to be further characterised. Indeed, a major challenge in gut microbiome research is linking the microbiota to specific physiological and metabolic responses. Promising developments such as the fish-gut-on-chip may offer opportunities to test the response of the fish gut microbiota to environmental parameters while studying the microbial populations *in vitro* [108]. The logical progression of this work would be to undertake a full metabolomics evaluation to provide a more robust interpretation of the significance of changes in the context of the functionality and whole animal response to variations in diet and the environment.

## Conclusions

Limiting the environmental impact of food production is paramount in ensuring future food security. We conclude that *Ulva rigida* species of macroalgae represent a low-cost, nutritious and palatable supplement to the diet of juvenile Atlantic cod (*Gadus morhua*). *U. rigida* can be supplemented up to 10% inclusion levels without negative impacts on the growth, gut morphology, or microbiome. We did not see any apparent increase in growth performance, though it should be noted that functional feed supplements and additives rarely manifest directly as having significant effects on improving fish growth rates under optimum rearing conditions. In contrast, we also determined that 10% dietary inclusion of *Ascophyllum nodosum* was not a suitable supplement for cod. This was evidenced by poor fish growth performance and had negatively impacted on gut integrity and appeared to negatively impact gut microbiome development and microbiome diversity. In the quest for economic and environmentally sustainable marine ingredients for aquafeeds, we advocate the potential for macroalgae (*U. rigida* in particular) to play a prominent role and support fish health and welfare as a sustainable feed ingredient.

## Supporting information

Supplementary information Doc

## List of Abbreviations

ICES: International Council for the Exploration of the Sea
CTRL: Control diet treatment
ULVA: *Ulva rigida* supplement diet treatment
ASCO: *Ascophyllum nodosum* supplement diet treatment
ASCO_N: *Ascophyllum nodosum* supplement diet treatment – Normal growth
ASCO_LG: *Ascophyllum nodosum* supplement diet treatment – Lower growth
LM: Light Microscopy
SEM: Scanning electron microscopy (SEM)
MV: Microvilli
AU: Arbitrary units
PBS: Phosphate buffer saline
CTAB: Cetyl trimethylammonium bromide
DNA: Deoxyribonucleic acid
RNA: Ribonucleic Acid
PCR: Polymerase chain reaction
OTU: Operational Taxonomic Unit
K: Fulton’s condition factor
ANOVA: Analysis of Variance
NMDS: Non-metric multidimensional scaling
PCoA: Principal Coordinates Analysis
sPLS-DA: Sparse Projection to Latent Structure – Discriminant Analysis
TSS + CLR: Total Sum Scaling followed by Centralised Log Ratio
ANF: Antinutritional Factor

## Declarations

### Ethics approval and consent to participate

The feeding trial was carried out at the Carna Research Station, Carna, Co. Galway, Ireland, a Health Products Regulatory Authority (HPRA) licensed institution. All personnel involved in the feed trial work was Laboratory Animal Safety Trained Ireland (LAST-Ireland) with individual authorisation.

### Consent for publication

Not applicable.

### Availability of data and material

Raw sequences have been deposited in the NCBI sequence read archive (SRA) under bioproject number PRJNA636649. Sequencing analysis scripts can be found at http://www.tinyurl.com/JCBioinformatics with information provided by Dr. Umer Zeeshan Ijaz.

### Authors’ contributions

**CK** dissected the fish for microbiome sampling, DNA extraction optimisation and extractions on the gut content, bioinformatics, and statistical analysis (excluding tank population data) and data interpretation. **MBW** managed the feeding trial, coordinated sampling, data management and statistical analysis of tank populations. **JH** ran the feeding trial, assisted with sampling for growth and carried out sampling for and analysis of data from light microscopy. **RD** ran the feeding trial, assisted with sampling for growth and carried out sampling for and analysis of data from SEM. **SW** ran the feeding trial, carried out sampling for growth and assisted with all dissections for microbiome, light microscopy, and SEM analysis. **AHLW** seaweed identification and collection designed the test diets and feed analysis. **SJD** designed the research study, supervised the growth and gut morphology data analysis and data interpretation. **RF** designed research study, EIRCOD Project Co-ordinator. **UZI** wrote the analysis scripts to generate the microbiota figures in this paper and performed the microbiota bioinformatic and statistical analysis with **CK**. **CJS** designed the research study and supervised the microbiome work. The study was designed by SJD, MBW, AHLW, CJS and RF. CK, MBW, AHLW, SJD, UZI and CJS wrote the manuscript, which was reviewed and edited by all authors. **RF** was EIRCOD Project Coordinator. All authors made a substantial contribution towards the research study and all authors (except RF) approved the final manuscript.

## Acknowledgements

We dedicate this paper to Richard Fitzgerald, a dear colleague and friend, 1957-2016. The authors also kindly acknowledge all staff at the Carna Research Station, Carna, Co. Galway.

