## Supplementary information Doc for "Temporal changes in the gut microbiota in farmed Atlantic cod (*Gadus morhua*) outweigh the response to diet supplementation with macroalgae": AM_Supplementary Information.docx

### Supplementary Material

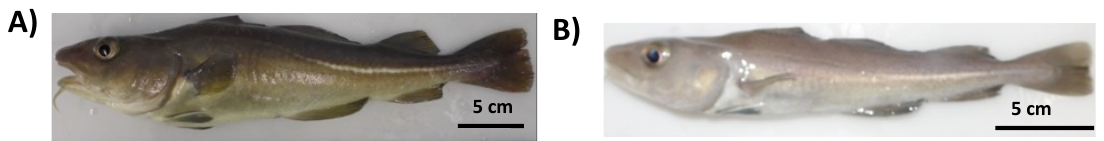

Figure S1. Photographic images of individual juvenile Atlantic cod (*Gadus morhua*) showing the visual distinction between A) ASCO_N, and B) ASCO_LG groups at Week 8. Note the smaller body size in the ASCO_LG individual which is consistent with poor food uptake.

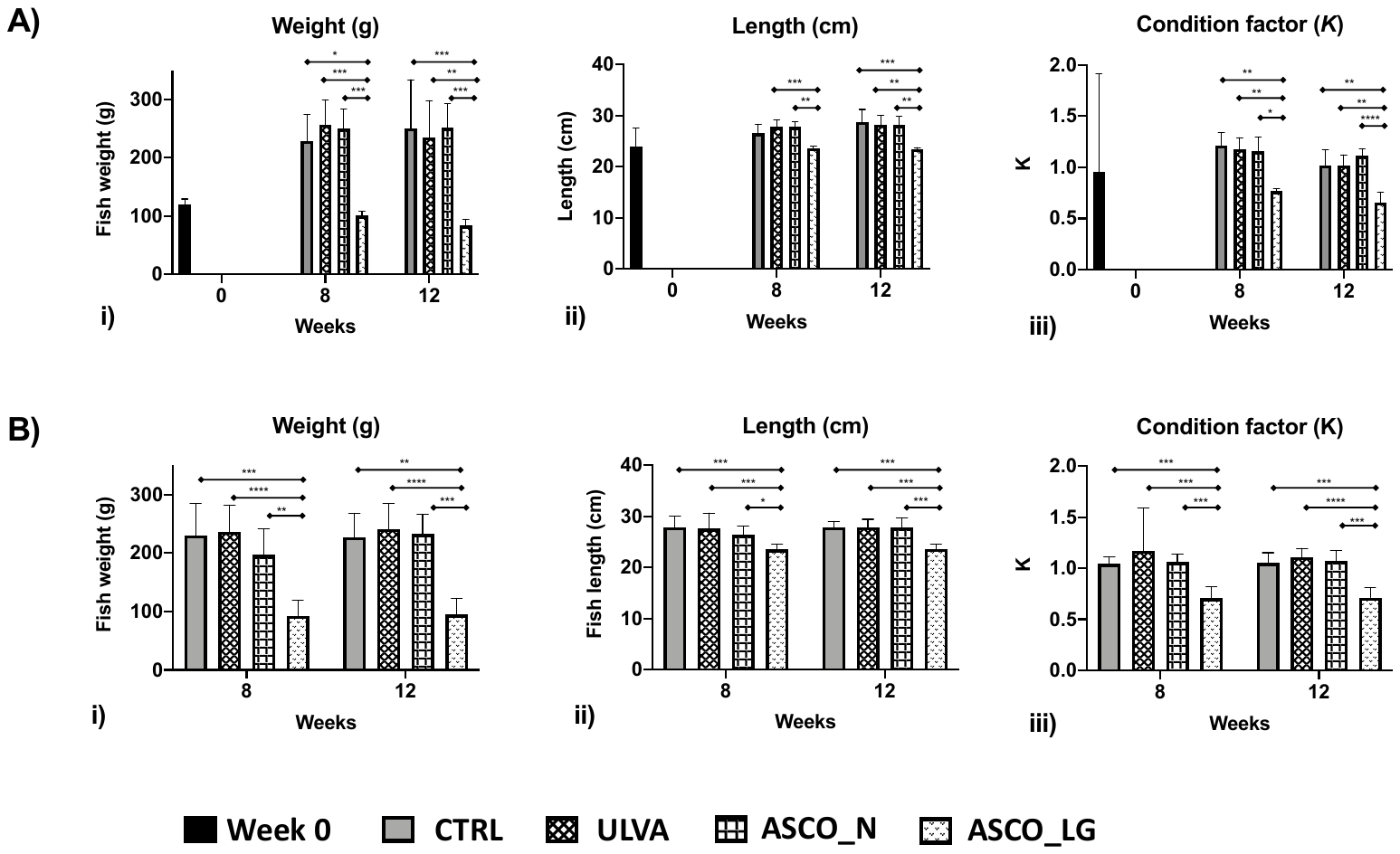

Figure S2. Bar-plots of fish sampled for microbiome analysis (A) and fish sampled for histology B) with i) the recorded weights (g), the recorded lengths (cm) in ii) and iii) the calculated condition factor K over the course of the trial. The associated statistical measurements were calculated using Kruskal-wallis significance testing.

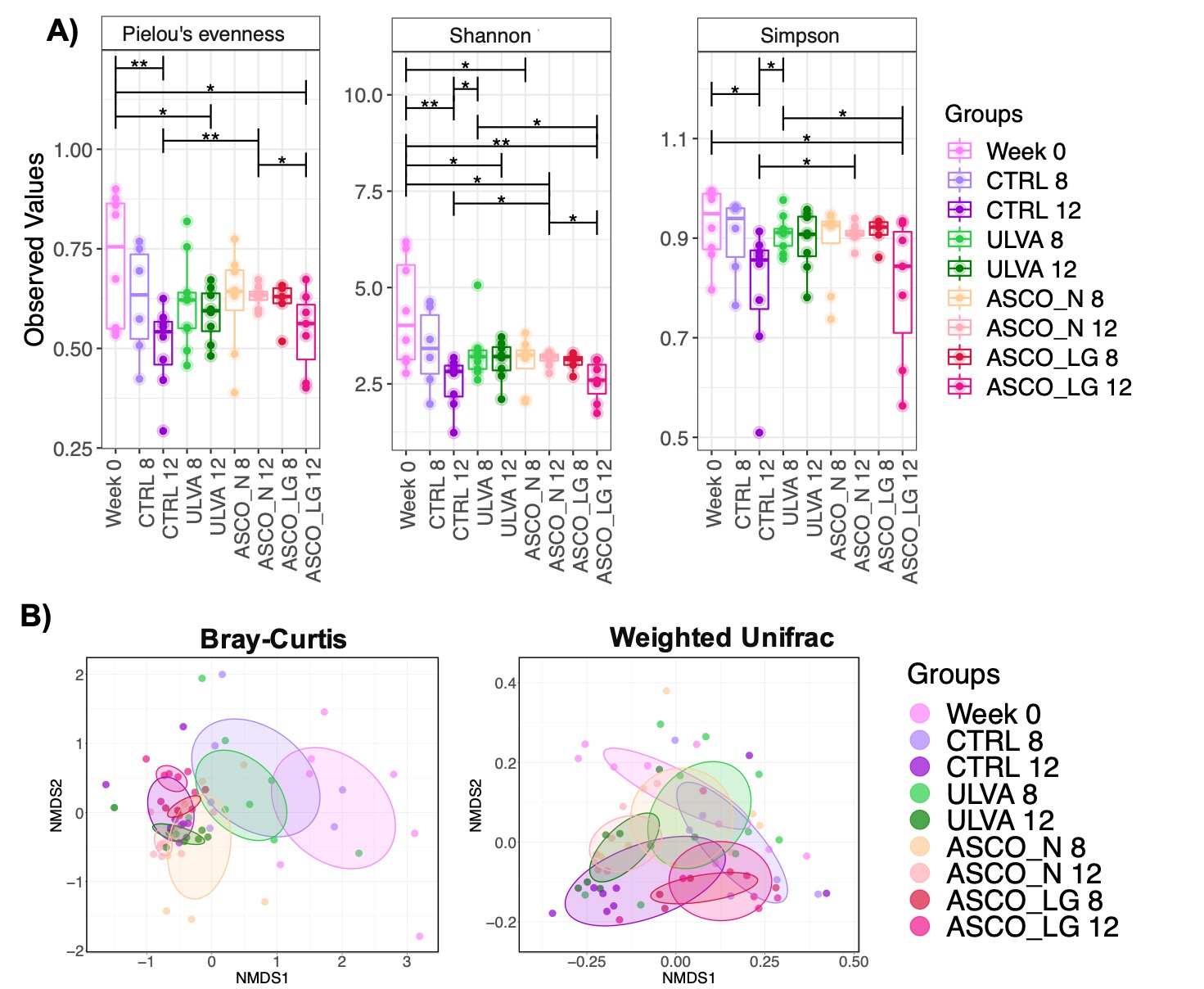

**Figure S3.** Microbial diversity and community structure according to variances in the 16S rRNA gene in DNA from 67 samples from Atlantic cod (*Gadus morhua*) microbiome. **A)** Alpha diversity box plots of Pielou’s evenness, Shannon Entropy and Simpson Diversity index. Pair-wise ANOVA *P*-values are displayed *P* < 0.05*, *P* < 0.01**, *P* < 0.001***. **B)** Beta diversity Non-Metric Multidimensional Scaling (NMDS) plots using Bray-Curtis dissimilarity (R^2^ = 0.24142, *P* = 0.001) and weighted UniFrac distances (R^2^ = 0.296, *P* = 0.001)., where each point corresponds to the community structure of one sample, groups are indicated by colour coded circles, the ellipses are drawn at a 95% standard error.

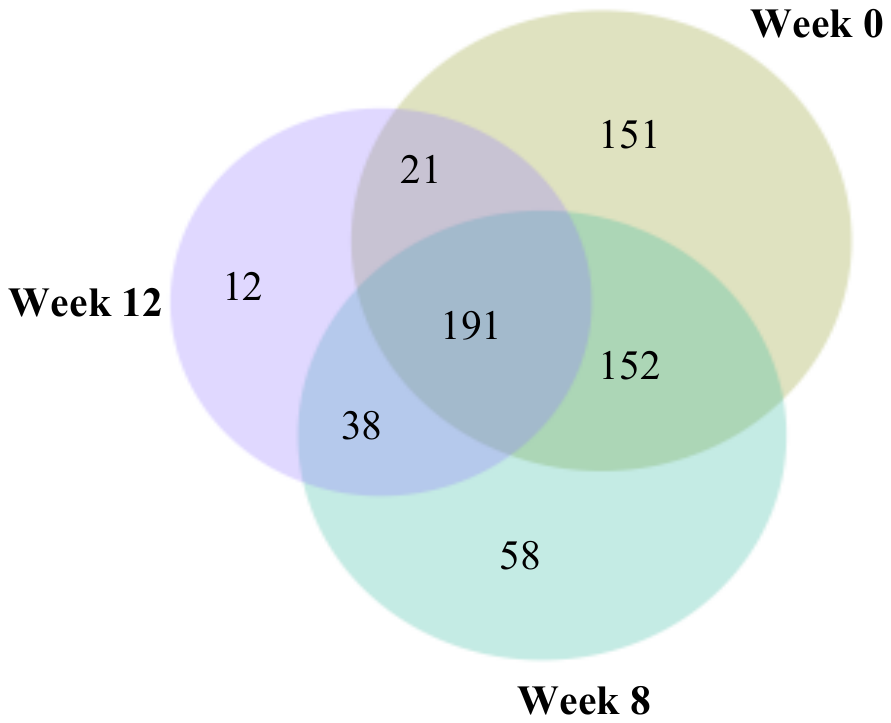

**Figure S4.** Venn diagram showing the number of shared operational taxonomic units (OTUs) and unique OTUs in the hindgut of juvenile Atlantic cod (*Gadus morhua*) from Week 0, Week 8 (CTRL, ULVA, ASCO_N and ASCO_LG and Week 12 (CTRL, ULVA, ASCO_N and ASCO_LG).

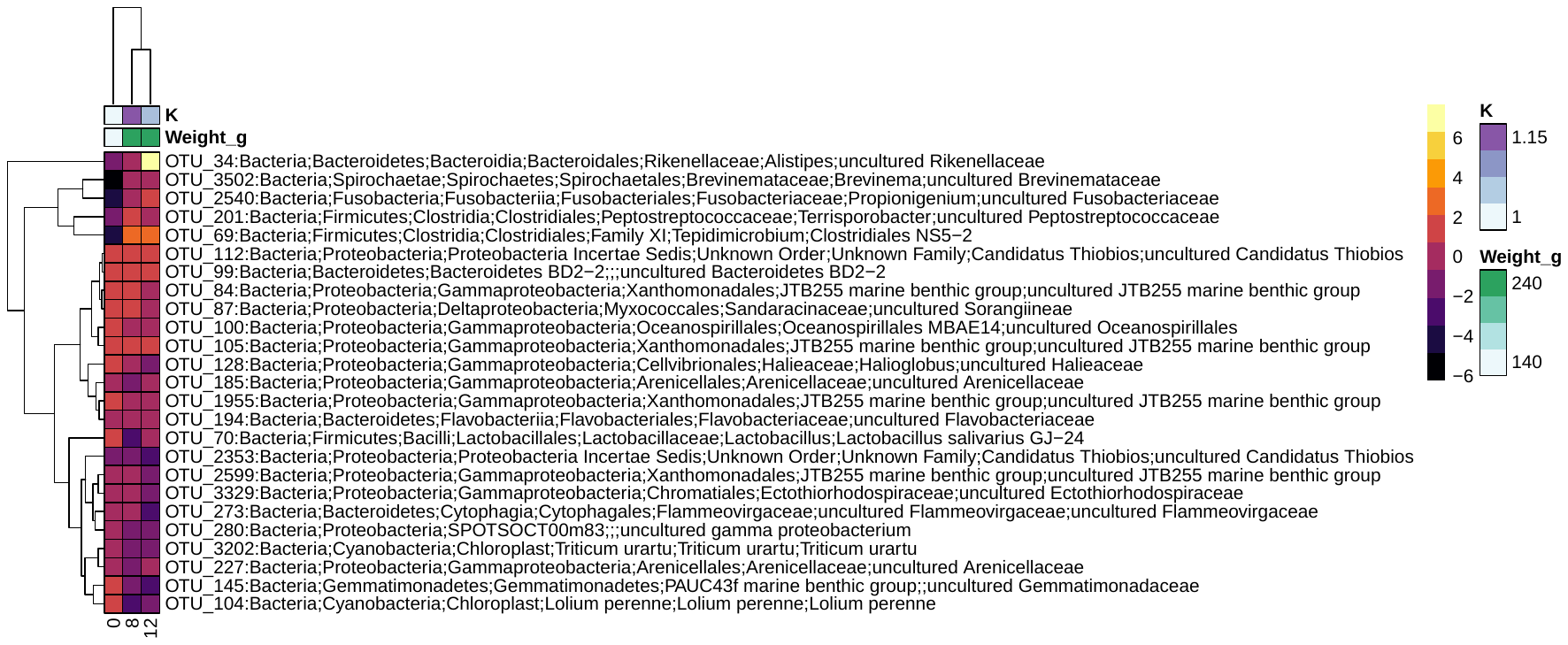

**Figure S5.** sPLS-DA heatmap of the discriminant operational taxonomic units (OTUs) > 1% relative abundance in the hindgut microbiota between the Week 0, Week 8 (CRTL, ULVA, ASCO_N and ASCO_LG) and Week 12 (CRTL, ULVA, ASCO_N and ASCO_LG) groups of juvenile Atlantic cod (*Gadus morhua*). Rows and columns are ordered using hierarchical (average linkage) clustering to identify blocks of genera of interest. The heatmap depicts TSS+CLR (Total Sum Scaling followed by Centralised Log Ratio) normalised abundances: high abundance (orange/yellow) and low abundance (dark purple). Fish weight (g) and condition factor K is shown per grouping. sPLS-DA was fine-tuned using centroids.dist and 3 tuning components.

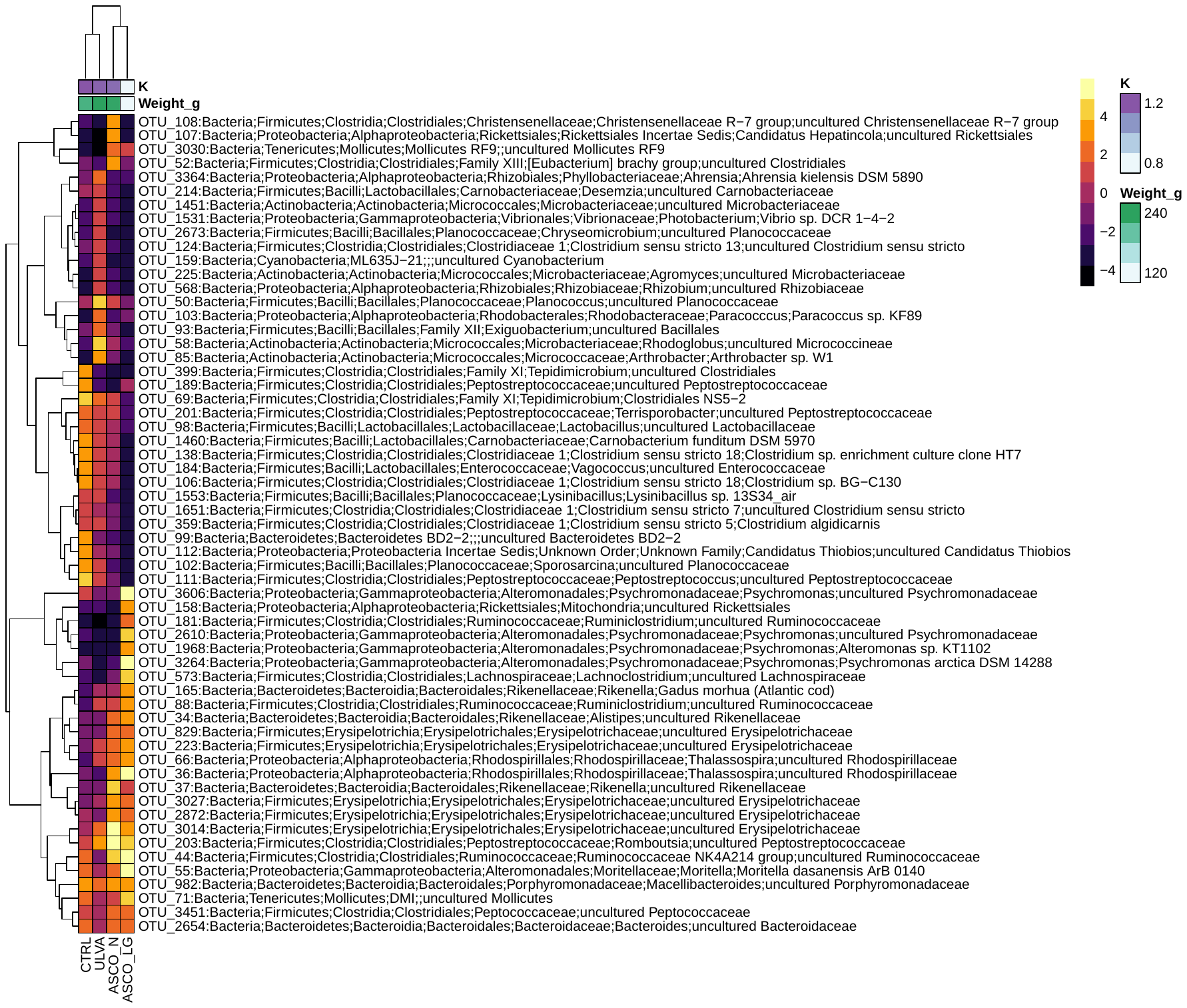

**Figure S6.** sPLS-DA heatmap of the discriminant operational taxonomic units (OTUs) > 3.5% relative abundance between the CRTL, ULVA, ASCO_N and ASCO_LG groups of juvenile Atlantic cod (*Gadus morhua*) at Week 8. Rows and columns are ordered using hierarchical (average linkage) clustering to identify blocks of genera of interest. The heatmap depicts TSS+CLR (Total Sum Scaling followed by Centralised Log Ratio) normalised abundances: high abundance (orange/yellow) and low abundance (dark purple). Fish weight (g) and condition factor K is shown per grouping. sPLS-DA was fine-tuned using centroids.dist and 3 tuning components.

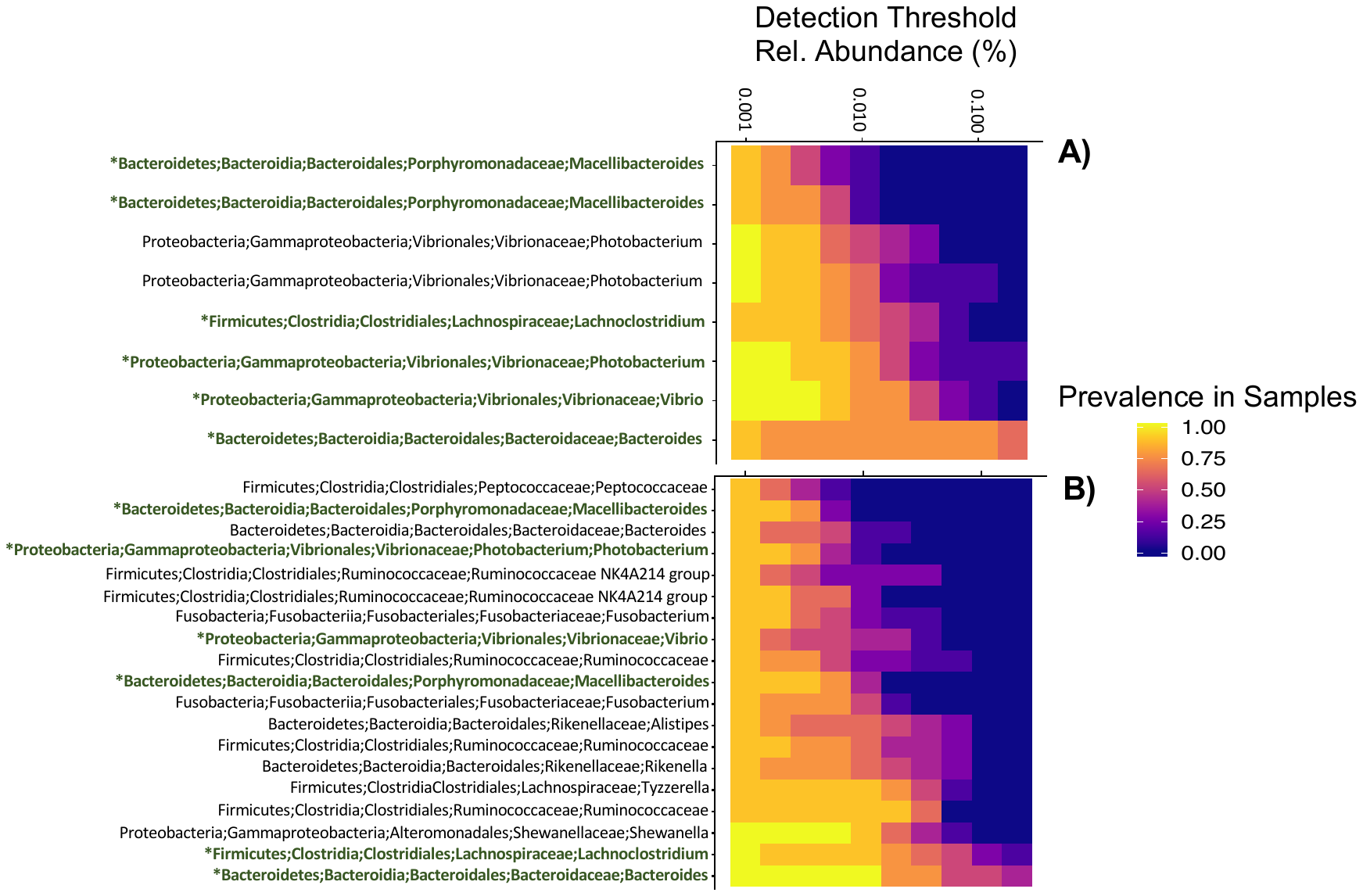

**Figure S7.** The core microbiome consisting of the operational taxonomic units (OTUs) found in 85% of individuals in the **A**) CTRL and **B**) ULVA dietary treatment at Week 12. Green text indicates those that are shared between the two dietary groups.
